# Integrated role of microRNA-30e-5p through targeting negative regulators of innate immune pathways during HBV infection and SLE

**DOI:** 10.1101/2020.02.29.969014

**Authors:** Richa Mishra, Sanjana Bhattacharya, Bhupendra S Rawat, Ashish Kumar, Akhilesh Kumar, Kavita Niraj, Ajit Chande, Puneet Gandhi, Dheeraj Khetan, Amita Aggarwal, Seiichi Sato, Prafullakumar Tailor, Akinori Takaoka, Himanshu Kumar

## Abstract

Precise regulation of innate immunity is crucial for the development of appropriate host immunity against microbial infections and the maintenance of immune homeostasis. The microRNAs are small non-coding RNA, post-transcriptional regulator of multiple genes and act as a rheostat for protein expression. Here, we identified microRNA(miR)-30e-5p (miR-30e) induced by the hepatitis B virus (HBV) and other viruses that act as a master regulator for innate immune responses. Moreover, pegylated type I interferons treatment to HBV patients for viral reduction also reduces the miRNA. Additionally, we have also shown the immuno-pathological effects of miR-30e in systemic lupus erythematous (SLE) patients and SLE mouse model. Mechanistically, the miR-30e targets multiple negative regulators namely *TRIM38, TANK, ATG5, ATG12, BECN1, SOCS1, SOCS3* of innate immune signaling pathways and enhances innate immune responses. Furthermore, sequestering of endogenous miR-30e in PBMCs of SLE patients and SLE mouse model respectively by the introduction of antagomir and locked nucleic acid based inhibitor significantly reduces type I interferon and pro-inflammatory cytokines. Collectively, our study demonstrates the novel role of miR-30e in innate immunity and its prognostic and therapeutic potential in infectious and autoimmune diseases.

## Introduction

The host innate immunity is an evolutionarily conserved defense system against microbial threats. These microbes express signature molecule known as pathogen-associated molecular patterns (PAMPs) which are sensed by host’s effectively conserved sensors known as pattern-recognition receptors (PRRs) in various compartments of cells. The coordinated interactions among them, activate a complex cascade of signaling pathways resulting in the development of innate immune responses for the elimination of invading microbes, through production of pro-inflammatory cytokines, type I and III interferons (IFNs) and chemokines, recruitment of immune cells and trigger various types of cell death^1^. These PRRs also interact with some host endogenous molecules known as danger-associated molecular patterns (DAMPs) and initiate innate immune responses without microbial infection and may establish or trigger autoimmune diseases^2^.

The micro RNA (miRNA) is a class of small non-coding RNA (18-22 nucleotide) that fine-tune protein expression through direct interaction with 3’UTR of the gene transcript^3^. miRNA interacts with target transcript through base pairing and initiates degradation or blocking of translation machinery via multiprotein complex known as RNA-induced silencing complex (RISC)^4^. It has been reported that a single miRNA has multiple mRNA targets and regulate cell signaling cascades and cellular responses during viral infections^5^. Several viruses evade immunity and establish infection by perturbing the host cellular miRNA expression or expressing viral(v)-miRNA upon infection^6^. In contrast, it has been reported that several host miRNAs restrict viral replication by targeting viral genome or those host genes which are essential for viral replication^7, 30^. In this study, we have identified miR-30e induced by DNA and RNA viruses such as Hepatitis B virus (HBV), Human cytomegalovirus (HCMV), New Castle disease virus (NDV) and Sendai virus (SeV) in primary cells such as human PBMCs and various mammalian cell lines. Notably, higher miR-30e levels were also detected in the serum of therapy naive HBV patients. Introduction of miR-30e into the cells promote production of pro-inflammatory cytokines, type I/III IFNs and globally enhances the innate immunity and therefore reduces viral load. However, HBV patients treated with pegylated type I IFNs present reduced HBV infection in terms of viral load and disease pathology, and significant concomitant reduction in miRNA30e levels, illuminating the correlation of innate immune responses and miR-30e expression. We utilized bioinformatic tools and transcriptomic approaches to identify miR-30e targets negative regulators namely *TRIM38, TANK, ATG5, ATG12, BECN1, SOCS1* and *SOCS3* of PRRs-mediated signaling pathways such as TLR, RLR and DNA sensing innate immune signaling pathways^8, 9^. Additionally, we showed miR-30e and 3’UTR of negative regulator transcripts makes a complex with Argonaute 2 (Ago2) protein, a component of RNA induced silencing complex (RISC), reduces transcript levels and subsequently protein expression thereby enhances innate immune responses. In contrast, patients with Systemic Lupus Erythematosus (SLE), a disease characterized by Type I IFN signature had higher expression of miR-30e^10, 11^. Further, NZB/NZW F1 hybrid, an animal model of lupus^31^ also showed increased expression of miR-30e. In connection, the introduction of miR-30e antagomir into PBMCs of SLE patients and lock nucleic acid (LNA) inhibitor for miR-30e through intra-orbital injection to the SLE mouse model significantly reduces type I IFNs and proinflammatory cytokines and moderately enhances the innate negative regulation. In conclusion, our study, proposes the novel role of miR-30e in innate immunity and its prognostic and therapeutic potential in HBV and SLE.

## Results

### miRNA-30e is induced upon viral infection and enhances innate antiviral responses to inhibit viral replication

To investigate the miRNAs involved in regulation of innate immune response during viral infections, we performed unbiased data analyses on previously published reports and miRNA microarray GEO datasets as shown in schematic workflow (Fig. S1A). In particular, the miRNA reports in H5N1 or Epstein Barr virus were analyzed for upregulated miRNAs. These upregulated miRNAs were compared to our previous miRNA profiling dataset from NDV infection in HEK293T cells (GSE65694). Upon comparison with NDV infection we selected miR-30e-5p, miR-27a-3p and mir-181a/2-3p as the common miRNAs across miRNA profiles related to viral diseases (Fig. S1B). Our analysis identified miR-30e as a unique miRNA that was predicted to target various PRR-mediated signaling regulators during negative regulation of innate immune responses (Fig. S1C) and was upregulated in viral infections moreover its mature form was highly conserved among the wide range of species (Fig. S1D). Additionally, datasets for H1N1 infection in mice (GSE69944), H5N1 infection in human lung carcinoma cells (A549 cells, GSE96857) and HBV infected liver tissues of hepatitis patients (GSE21279) were also analyzed by GEO2R package for upregulated miRNAs and among all upregulated miRNAs, miR-30e upregulation is represented here (Fig. S1E). The expression of miR-30e was upregulated during viral infections or stimulation with PAMPs *in-vitro* in various cell-lines (Fig. S2A-F). At the transcriptional level, miR-30e promoter activity was moderately enhanced by NDV but was unaffected by *IFNβ* or *TNFα* stimulation which activated the *ISRE* and *NFκB* promoters respectively, suggesting that miR-30e expression might be induced by the viral infections but not by the cytokines produced during infection (Fig. S2G-L). To understand the clinical relevance, induction of miR-30e was tested in the cohort of 51 non-treated HBV patients (demographic details mentioned in Table T1). To this end, the expression of miR-30e was evaluated from serum samples of therapy naive chronic hepatitis B (CHB) patients in comparison with healthy controls, and significantly elevated levels of miR-30e were detected in HBV patients (Fig. 1A). Similar results were obtained with HepG2 cell line treated with serum from HBV patients (HBV PS) for different time points, as shown (Fig. 1B), the induction of miR-30e enhanced significantly at 1*dpi*, 2*dpi* and 3*dpi* (days post infection) with maximum expression at 3*dpi*. Additionally, miR-30e mimic (miR-30e) inhibited HBV infection in HepG2 cells treated with serum from HBV patients or HepG2215 cells, stably expressing HBV replicon cells as compared to control miRNA (miR-NC1) treated cells as tested by HBV-specific RNA and DNA (Fig. 1C and D) (Fig. S3A). Interestingly, ectopic expression of miR-30e significantly reduced the HBV replication in HepG2-NTCP cells as tested by HBV RNAs and HBV pgRNA (Fig. S3B). Notably, we found significant elevated levels of HBV DNA and HBV covalently closed circular DNA in HBV patient serum infected HepG2 cells and stably expressing HBV replicon HepG2215 cells compared to uninfected HepG2 cells (Fig S3C) to show the infection in cells. On another hand, expression of *IFNλ1* and *IFIT1* was enhanced in HepG2 and HepG2215 cells, respectively, in the presence of miR-30e (Fig. 1C and D) (Fig. S4A). Similarly, HBV infection in HepG2-NTCP cells (liver hepatoma cell line permissive for HBV infection) overexpressing miR-30e (ectopic), elevated IFN*λ*1 transcript (Fig. S4B). To study whether miR-30e was involved in controlling RNA virus infection, we infected human PBMCs (hPBMCs) with NDV to quantify the expression of miR-30e. We found that NDV infection elevated the expression of miR-30e in time-dependent manner (Fig. 1E). Additionally, PBMCs infected with NDV in presence of miR-30e showed a significant reduction in NDV replication with a concomitant elevation of *IL6* expression whereas miR-30e inhibitor (AmiR-30e) reversed this phenomenon (Fig. 1F). Similar inhibition of viral replication was observed in multiple cell lines infected with NDV in the presence of miR-30e or AmiR-30e (Fig. S3D-F). Comparable results for antiviral responses were obtained after NDV infection in different cell-types at transcript level and protein levels (Fig. S4C-I) in presences of miR-30e and it also activate *ISRE, IFNβ* and *NFκB* promoters as tested by luciferase assay (Fig. S4J-L). Furthermore, miR-30e presence reduces NDV replication in terms NDV protein as tested by NDV-specific antibody estimated by western blot, microscopy and FACS analysis respectively (Fig. 1G-H and S3G). To further validate the function of miR-30e in controlling viral infections, we quantified the expression of miR-30e upon Sendai virus (SeV) infection in A549 cells. SeV induced the expression of miR-30e (Fig.1I) and ectopic expression of miR-30e (miR-30e mimic) inhibited the SeV replication as shown by qRT-PCR and FACS analysis (Fig. 1J-K) through enhancing antiviral genes such as the expression of *IFIT1* (Fig. 1J).

**Figure 1.**
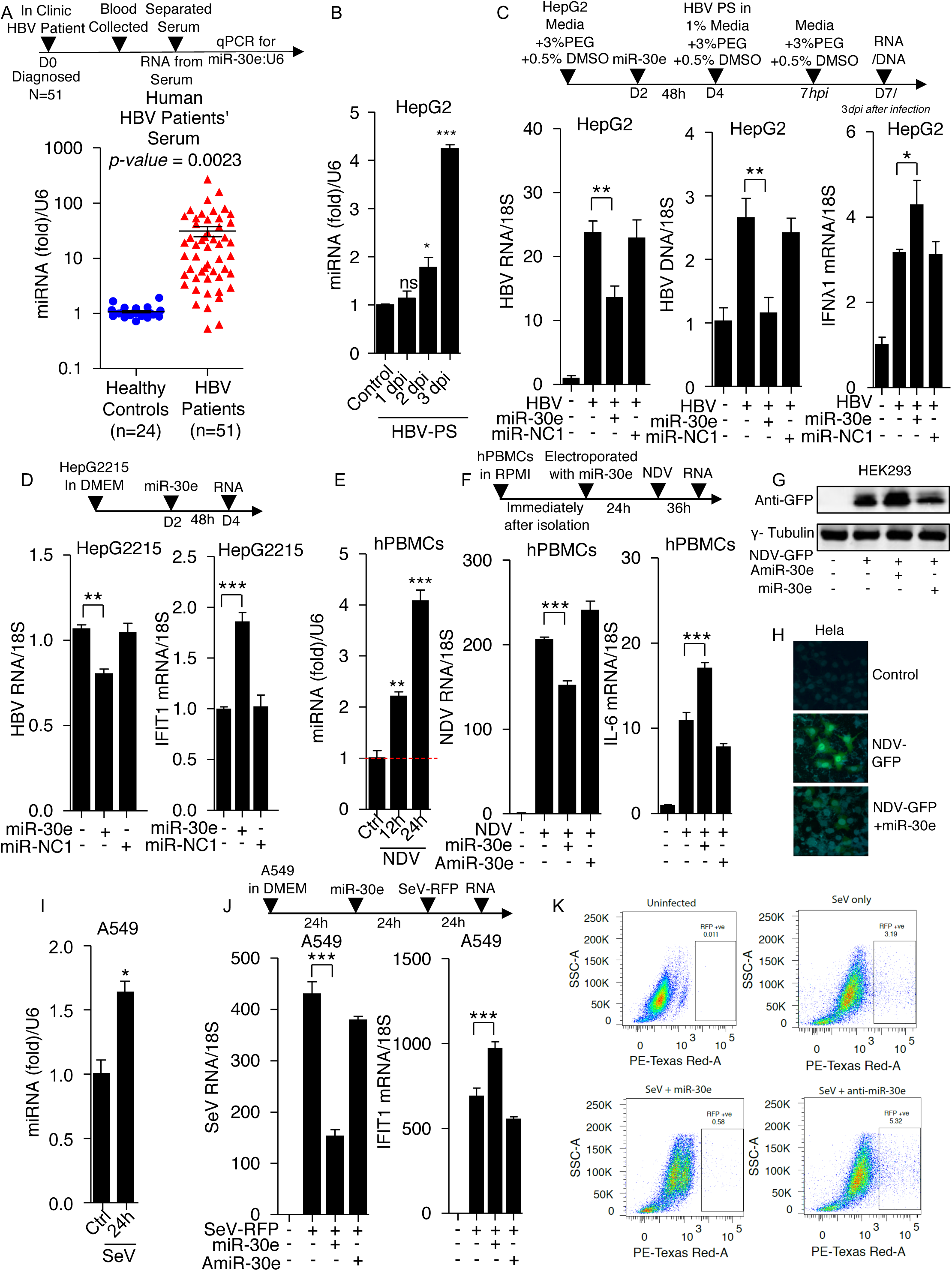
Viral infection induces miRNA-30e that inhibits virus replication by promoting innate immunity. (A) Quantification (as determined by qRT-PCR analysis) of the fold changes in the abundances of miR-30e as indicated, in the serum collected from hepatitis B patients (n=51) compared to healthy controls (n=24). (B-F; I and J) Quantification of the fold changes in the relative abundances of miR-30e, viral transcripts and respective innate immune transcripts (*IFNλ1, IL6 and IFIT1)* at the indicated times after treatment or infection with (B) HBV patient’s serum (HBV PS) in HepG2 cells, (C) HepG2 cells were transfected with miR-30e (50 nM) or miR-NC1 (50 nM) prior to infection (D) HepG2215 cells, stably expressing HBV replicon HepG2 cells transfected with miR-30e or miR-NC1, (E) NDV (MOI 5) in human PBMCs, (F) hPBMCs transfected with miR-30e or AmiR-30e (50 nM) prior to infection and (I) SeV (MOI 5) in A549 cells (J) A549 cells transfected with miR-30e or AmiR-30e prior to infection. (G, H and K) Quantification of viral infection as indicated in (G) HEK293 cells (transfected with miR-30e or AmiR-30e for 24 hours then infected with GFP-tagged NDV (NDV-GFP) (MOI 5) for 36 hours and subjected to immunoblot analysis using antibodies specific for GFP (anti-GFP antibody) and *γ*-tubulin (used as a loading control), (H) HeLa cells transfected with miR-30e or infected with NDV-GFP and subsequently subjected to confocal microscopic analysis for NDV particles with anti-GFP antibody (green) and, nuclei were visualized with 4′,6-diamidino-2-phenylindole (DAPI; blue) and (K) A549 cells were transfected with miR-30e or AmiR-30e for 24 hours then infected with RFP tagged SeV (MOI 5) for 24 hours and analyzed by flow cytometery. Ctrl represents control untreated sample, D; represents number of days, *dpi*; represents days post infection and *hpi*; represents hours post infection. Data are mean +/-SEM of triplicate samples from single experiment and are representative of three (A-C, E, F, I, J) two (D, G, H, K) independent experiments. \*\*\**P<*0.001, \*\**P<*0.01 and \**P<*0.05 by one-way ANOVA Tukey test, Mann-Whitney test and unpaired t-test.

Next, role of miR-30e on DNA virus replication was determined, to this end, HFF cells were infected with DNA virus, HCMV, alone or along with miR-30e or miR-NC1. The viral replication was significantly reduced in the presence of miR-30e compared to the miR-NC1 treated HFF cells as quantified by HCMV transcript encoding viral glycoprotein gene (Gly B) by real-time PCR and analyzed by microscopy (Fig. S5A) and additionally the transcript levels of *IL-6* was enhanced (Fig. S5B).

To investigate, how miR-30e influence antiviral responses upon treatment with pure viral ligand such as poly IC and ssRNA, different cells including human PBMCs were stimulated along with miR-30e.The expression of various cytokines and ISGs such as *IFNβ, IP-10, IL-6* and *IFIT1* were elevated upon poly IC or ssRNA stimulation along with miR-30e transfection and similar results were found for the promoter activity of *ISRE* and *IFNβ* in presence of miR-30e (Fig. S6A-H). Collectively, our results demonstrate that miR-30e is upregulated in during virus infection and miR-30e inhibits viral replication by promoting the expression of innate antiviral genes.

### miR-30e globally enhances innate immune responses during virus infection

To investigate the effect of miR-30e on innate immune responses interms of innate immune immune genes upon virus infection, A549 cells were either mock transfected, or transfected with miR-NC1 and miR-30e for 24 hours, followed by infection with NDV for 24 hours and finally subjected to whole transcriptome sequencing using an illumina next-generation sequencer (NGS) and analyzed for differentially expressed genes as shown in the schematic (Fig. 2A, S7A). Notably, the transfection efficiency of miR-30e in both replicates was confirmed by qRT-PCR using miR-30e-5p Taqman and reduction in viral infection was confirmed by quantifying the NDV RNA in both the replicates to establish further analysis as per our previous findings (Fig. 2B). Principal component analysis for the samples resulted in formation of three distinct groups (miR-30e, miR-NC1 and uninfected) according to their treatment (Fig. 2C). Additional analysis of transcriptomic data showed that 1179 genes were significantly upregulated and 1206 genes were significantly downregulated upon miR-30e transfection in comparison to miR-NC1 (Fig. 2D (represented in volcano plot) and Fig. S7B (represented in MA plot and shown in Table T2). Moreover, KEGG pathway analysis of significantly upregulated genes, upon miR-30e treatment and NDV infection indicated enrichment of genes belonging to key cellular mechanisms namely: cell cycle, NOD-like receptor signaling pathway, MAPK signaling pathway, TLR signaling pathway, RLR signaling pathway, PI3K-AKT signaling pathway, cytokine-cytokine receptor interaction pathway, NF-kappa B signaling pathways, TNF signaling pathway, etc. (Fig. 2E). Relative expression levels of top genes involved in these pathways is represented by heat map (Fig. S7C). Intriguingly, we noticed that a significant number of interferons stimulated genes (ISGs) like *IRF3, IRF7, CXCL10 (IP10), IFIT1, MX1, IL6, OAS1* among others were also predominantly upregulated (Fig. 2F) that confirmed with the initial findings of the study. Furthermore, our NGS results were verified by quantifying the expression level of type 1 interferon *IFNβ*, interferon stimulated genes: *IFIT1* and *OAS1* and pro-inflammatory cytokines: *IL6* and *IP10*. These outcomes intricate strongly that miR-30e reduces the viral replication by enhancing the innate immune responses upon activation of various signaling cascades. Additionally, miR-30e impacts innate immune responses during viral infection, prompted us to investigate transcriptome and gain mechanistic insight for the target of miR-30e.

**Figure 2.**
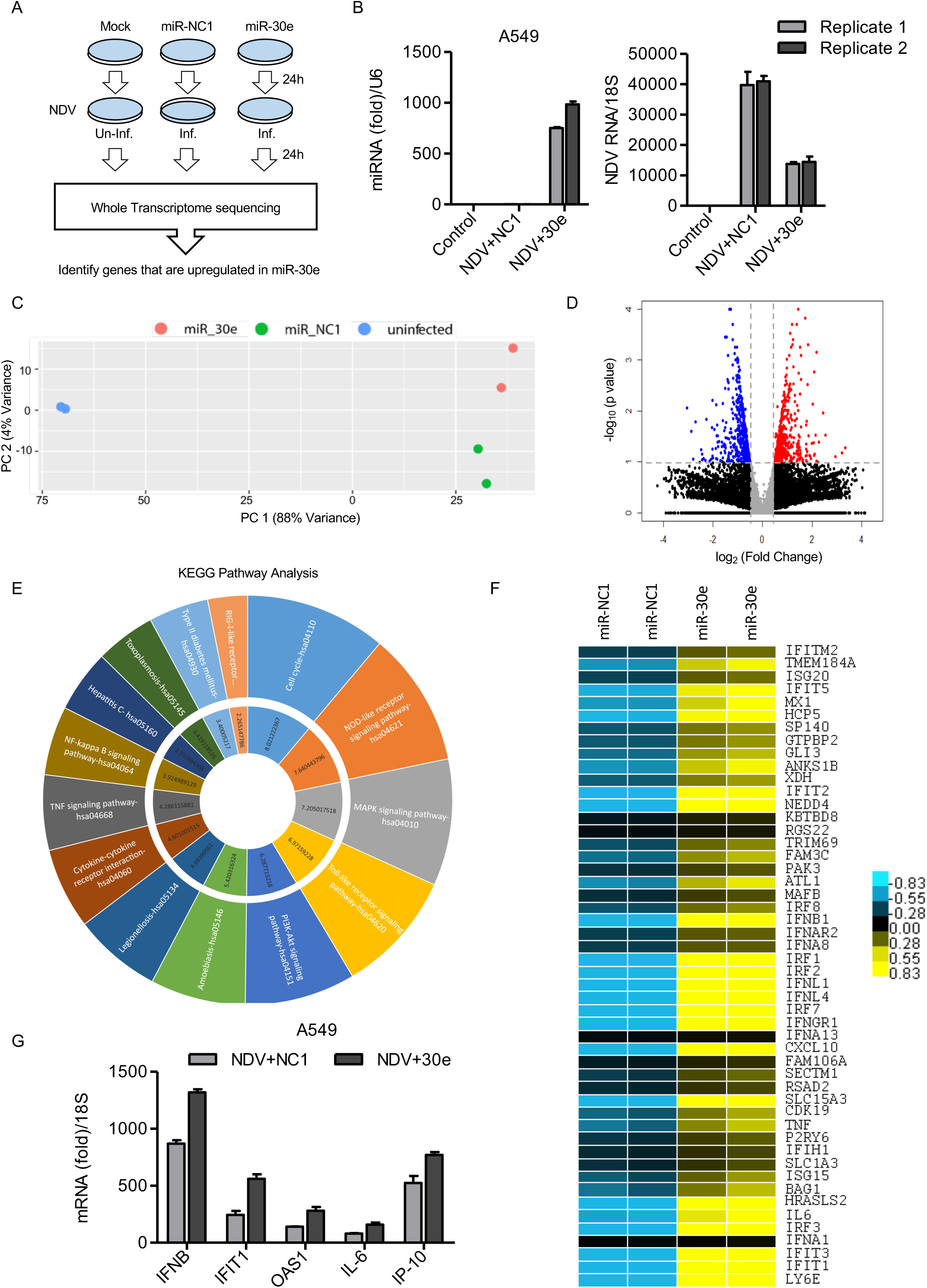
Transcriptomic analysis shows miR-30e enhances innate immune responses during NDV infection: (A) Schematic outline of transfection with control (miR-NC1) or miR-30e and NDV infection (MOI 5) in A549 cells at indicated time and subjected to whole transcriptome sequencing and gene analysis (Inf: infected and Un-inf: uninfected). (B) Quantification of the fold changes in the abundances of miR-30e is measured by qRT-PCR and normalized by U6 control and NDV viral transcripts in both the replicate samples used for transcriptome sequencing and analysis. (C) Plot showing first two components from principal component analysis of all the 6 samples, distance between samples indicate how different they are from each other in terms of gene expression. (D) Volcano plot represents differential expression of genes between two groups of samples (miR-30e and miR-NC1 overexpression) during NDV infection in A549 cells. For each gene: *P-value* is plotted against fold change (miR-30e vs miR-NC1). Genes significantly changed (>1.5-fold) are colored in red (upregulated) and blue (downregulated). (E) KEGG pathway analysis of upregulated genes, outer circle indicates top upregulated pathways and the inner circle represents corresponding combined score (a derivative of *P-value* and Z-score). (F) Heat map represents relative abundance of top upregulated interferon stimulated genes across different samples. (G) Quantification (measured by qRT-PCR) of the fold changes in the abundances of type 1 interferon and pro-inflammatory cytokines in the samples, A549 cells transfected with miR-NC1 or miR-30e and infected with NDV as indicated (NDV+NC1) and (NDV+30e), analyzed by RNA-Sequencing.

### miR-30e targets negative regulators of TLRs, RLRs, DNA sensing and interferon signaling pathways

To investigate underlying molecular mechanism for the reduction of viral burden and enhanced antiviral innate immune responses by miR-30e, we conducted unbiased rigorous screening using various bioinformatic tools for the identification of innate immune genes.

To filter the genes transcript targeted by miRNA, certain criteria for screening were applied. First of all, common genes involved in innate immune regulation upon viral infections and targeted by miR-30e were identified and subjected to KEGG pathway analysis. The analysis revealed that majority of genes (*TRIM38, TANK, ATG5, ATG12, BECN1, SOCS1, SOCS3, TRIM13* and *EPG5*) were involved in negative/down regulation of pattern recognition receptors (PRRs)-mediated signaling pathways (Fig. 3A). Additionally, our NGS analysis demonstrated that expression of identified negative regulators during NDV infection were reduced upon miR-30e treatment as compared to miR-NC1 group (Fig. 3B). This further concludes that negative regulators were targeted by mir-30e. The binding efficiency for miR-30e and identified targets genes are significantly high to alter any physiological functions by the miRNA that was tested by different *in-silico* tools such as miRanda, DIANA, targetscan, miRDB and BiBiServ2_RNAhybrid as reported (Table T3). Although few targets of miR-30e are not negative regulators however, binding energy for those target transcript and miRNA assembly are low (Fig. S8) and therefore, it may not significantly alter the cellular function. To test specificity of miRNA with the 3’UTR of identified negative regulator genes, the 3’UTR of the gene were cloned downstream of luciferase gene under the CMV promoter and performed luciferase assy. It was found that miR-30e significantly reduced luciferase activity of investigated genes compared to control miR (Fig. 3C). In contrast, introduction of mutation in cloned 3’UTR by site directed mutagenesis (SDM) did not change the luciferase activity in presence of miR-30e and it was comparable with miR-NC1 (Fig. S9A), moreover, it could be noted here that, the target sequence in each negative regulator genes were same. Additionally, we knockdown these negative regulators in HEK-293T cells and Infected them with NDV to further estimated the level of *IFNβ*, which clearly elucidated their inhibitory effect on the mRNA levels of *IFNβ* within a cell, this effect was found to be significant with respect to the majority of the targets (Fig. 3D). And observed that the production of *IFNβ* is comparable after knockdown of target genes and introduction of miR-30e into the cells during viral infection suggesting the pivotal role of miR-30e in suppression of negative regulators transcripts. Furthermore, we scanned the 3’UTRs of identified negative regulators for RNA binding site for Ago2 protein in CLIP database, which is a key component of the miRNA-mediated silencing complex (RISC) and found that the miR-30e strongly complexes with the target genes as shown in (Table T4). To validate, miR-30e and negative regulator transcripts *(TRIM38, TANK, ATG5, ATG12, BECN1, SOCS1 and SOCS3)* interaction, Ago2 pull-down assay was performed as shown in schematic diagram (Fig. 3E) and found that introduction of miR-30e significantly enriches the transcript of negative regulators compared to the NDV alone infection or NDV infection along with control miRNA treated cells suggesting that miRNA directly interact with the transcript through the formation of RISC.

**Figure 3.**
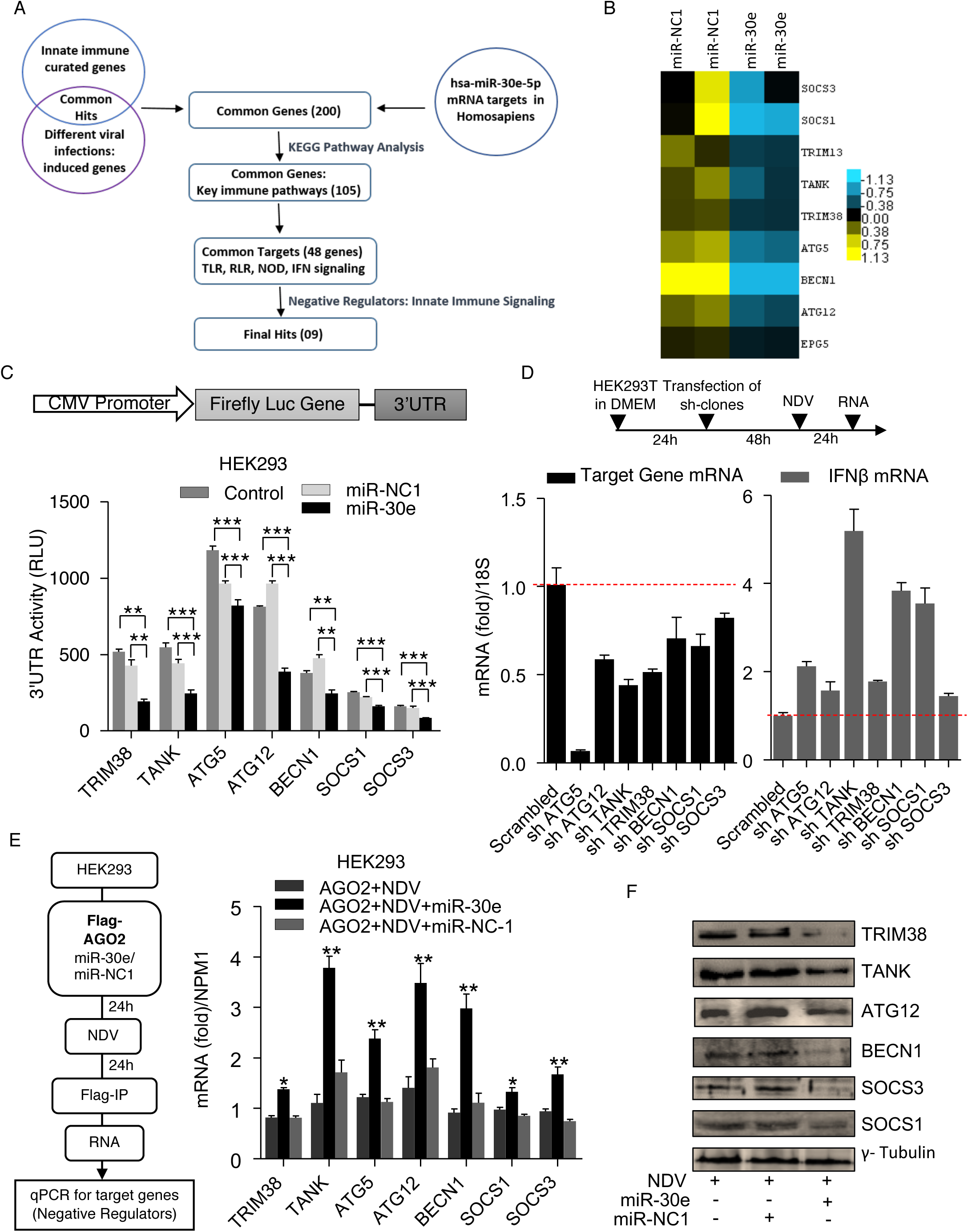
miR-30e targets 3’UTR of negative regulators of innate immune signaling pathways: (A) Screening pipeline used for identification of miR-30e target genes based on the indicated schematic workflow, final hits 09 genes corresponds to negative regulators targeted by miRNA-30e. (B) Reanalysis of previous transcriptome data for identification of negative regulators (targeted by miR-30e) upon miR-30e transfection and NDV infection (MOI 5) compared to miR-NC1 in two replicate samples. (C) HEK293 cells were transfected with 50 ng of pRL-TK and 300ng of 3’UTR_WT (of indicated genes) together with 25 nM miR-30e or miR-NC1, 24 hours after transfection, the cell was lysed and subjected to luciferase assay. (D) HEK293T cells were transiently transfected with 1.5μg of *sh-*clones of indicated genes or scrambled control for 48 hours then infected with NDV (MOI 5) for 24 hours and subjected to the quantification of the indicated transcripts or genes and *IFNβ*. (E) Schematic for RNA-immunoprecipitation assay. HEK293 cells were transfected with plasmid encoding Flag-Ago2 in presence of miR-30e (50 nM) and miR-NC1 (50 nM) and then infected with NDV (MOI 5). 24 hours after transfection cells were subjected to RNA immunoprecipitation with ant-Flag antibody and quantified for *TRIM38, TANK, ATG5, ATG12, BECN1, SOCS1* and *SOCS3* transcripts. (F) A549 cells were transfected with miR-30e or miR-NC1 mimic and then infected with NDV (MOI 5) for 36 hours before being subjected to immunoblot analysis with antibodies specific for indicated protein or *γ*-tubulin (used as a loading control). Data are mean +/-SEM of triplicate samples from single experiment and are representative of three (C) two (D, E, F) independent experiments. \*\*\**P<*0.001, \*\**P<*0.01 and \**P<*0.05 by one-way ANOVA Tukey test and unpaired t-test.

Next, the expression of identified negative regulators were examined in A549 cells upon NDV infection and found that at 12 hours there was increased in the expression of *TRIM38, TANK, ATG5, ATG12, BECN1, SOCS1* and *SOCS3* transcript and it was reduced at 24 hours due to induction of miR-30e as shown in left panel (Fig. S9B). Additionally, ectopic expression of miR-30e reduced the expression level of these targets in A549 cells compared to the control after NDV infection (Fig. S9C). Consistent with these results induction of targets was also reduced in HBV-patient serum treated HepG2 cells, HepG2215 cells (stably expressing HBV replicon cells and HepG2-NTCP cells infected with HBV, in presences of miR-30e compared to the control miRNA transfection (Fig. S10A-C), suggesting that miR-30e targets these genes during HBV and NDV infection in HepG2, HepG2215, A549 cells, respectively. Similar results were obtained due to miR-30e transfection in NDV infected and poly IC treated HeLa cells (Fig. S10D-E). We not only confirmed the expression of negative regulators by analyzing transcripts but also tested for protein expression using specific antibodies by immunoblot analysis. The introduction of miR-30e significantly reduced the expression of *TRIM38, TANK, ATG12, BECN1, SOCS3* and *SOCS1* as shown (Fig. 3F). Therefore, our results strongly suggest that these key negative regulators of innate immunity are targeted by miR-30e, which are induced during virus infection and resulting to enhanced antiviral responses.

### DAMPs induce miR-30e and enhance innate immune responses

Our observation for induction of miR-30e and subsequent heightened innate immune responses upon viral infection or pure PAMP stimulation prompted us to investigate the ability of host DAMPs for induction of miR-30e because sustained DAMPs production in the host can lead to enhance sterile inflammation and subsequently it may establish autoimmune disease^3^. To this end, ex-vivo experiment was performed using hPBMCs from three healthy volunteers. The genomic DNA were extracted from a portion of hPBMCs and sonicated to make small size (approx. 110-150 bps) for efficient transfection into the cells (Fig. 4A). The cultured hPBMCs were stimulated with the extracted small size DNA and tested for the induction of miR-30e and innate immune cytokines. The DAMP stimulation significantly induces the miR-30e and expression of *IFNα, IFNβ, IFIT1, IL6* genes in all 3 healthy volunteers (Fig. 4C-E). Next, to understand physiological significance of DAMP-induced innate immune cytokines, the DAMP-stimulated cells were infected by NDV and NDV replication was measured. The dsDNA-stimulation significantly reduced the NDV replication (Fig. 4F) suggesting that inflammatory cytokines and type I interferons induced by dsDNA inhibited the viral replication. Although inhibition of viral replication could be the collective results of both dsDNA and virus-mediated induction proinflammatory cytokines, type I interferons and type I inducible genes. Finally, we examined the levels of apoptosis induced by autologous dsDNA in hPBMCs by Annexin-PI assay using FACS analysis as shown in Fig. S11A, dsDNA stimulation enhanced apoptosis compared to the mock stimulation and it is comparable to the positive control treated cells, by Camptothecin. Additionally, the levels of *TLRs 3/7/9* was estimated upon dsDNA treatment as previously it has been reported that DAMPs enhance the levels of these *TLRs.* Consistent with previous observation, we obtained similar results (Fig. S11B-D). Taken together, these results showed that DAMPs/dsDNA enhances miR-30e, innate immune responses as well as promote apoptosis and induce TLRs, the crucial characteristic features for the development of autoimmune disorder in co-occurrence with DAMPs/dsDNA and miR-30e might play pivotal role in autoimmune disease.

**Figure 4.**
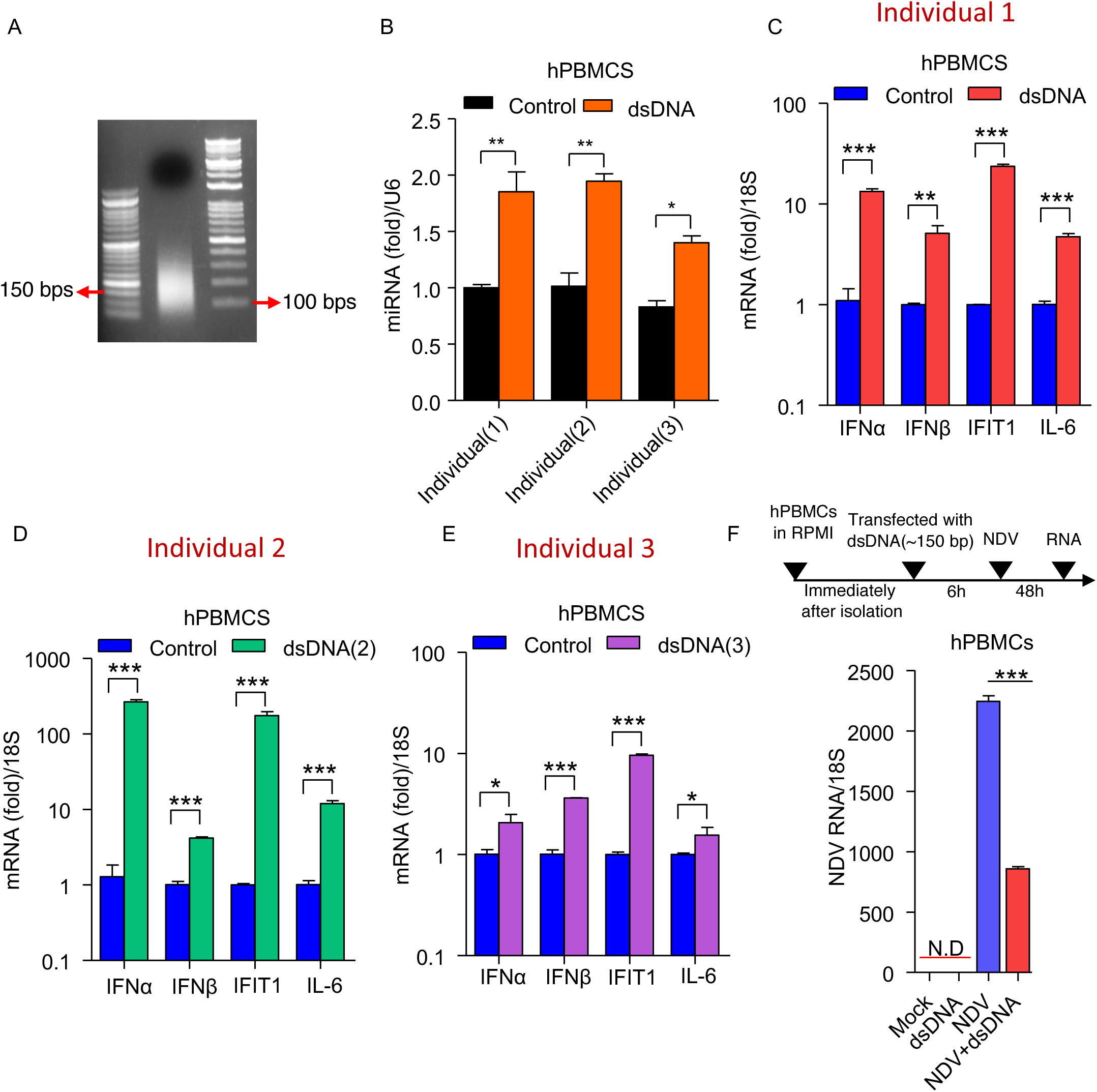
Human PBMCs stimulated with DAMPs induce miRNA-30e and enhance innate immune responses: PBMCs from three healthy individuals were transfected with their own genomic DNA (ds DNA). (A) Isolated genomic DNA sonicated into small fragments of dsDNA of approximately 100-150 bps each (as shown) and transfected using Lipofectamine 2000. (B-F) Quantification (by qRT-PCR analysis) of the fold changes in the relative abundances of (B) miR-30e, (C-E) respective innate immune transcripts (*IFNα, IFNβ, IFIT1* and *IL6)* in all individuals and (F) NDV viral transcript (shown as per schematic workflow). Data are mean +/-SEM of triplicate samples from single experiment and are representative of three independent experiments in three different individuals. \*\*\**P<*0.001, \*\**P<*0.01 and \**P<*0.05 by one-way ANOVA Tukey test and unpaired t-test, N.D. correspond to not detected.

### SLE patients and SLE mouse model show enhanced miR-30e expression

To investigate the role of miR-30e under physiological condition, an autoimmune disease, SLE was selected because SLE patients show enhanced inflammatory cytokines, type I interferons and type I interferon-inducible cytokines production^10^. The SLE patients also had elevated levels of several autoantibodies particularly antinuclear and anti-dsDNA antibodies. Therefore, PBMCs were isolated from clinically verified SLE patients (P: n=13) as shown (Table T5) and healthy controls (HC: n=13) and cultured for 48 hours, the expression levels of *IFNβ, IFIT1 and IL6* were compared by qRT-PCR. As expected, we found that the expression levels of *IFNβ, IFIT1 and IL6* were significantly enhanced in SLE patients compared to healthy controls (Fig. 5A). The enhanced innate immune responses prompted us to investigate the expression levels of miR-30e. Interestingly, the expression of miR-30e is significantly enhanced (several fold) in patients (n=13) compared to healthy controls (n=13). Further to confirm our observations in SLE patients, the SLE mouse model was used. The New Zealand white/black (NZW/B) mice were extensively used for SLE studies. The splenocytes from both parents NZB and/or NZW mice (n=7) mice and lupus prone F1 (F1: NZW/B n=7) generation mice were tested for the expression levels of *Ip10, Tnfα and Il6* by qRT-PCR (Fig. 5B). The F1 mice showed significantly high level of inflammatory responses compared to non-SLE parent mice. Consistent with human SLE results, the expression of miR-30e is enhanced manifold both in F1 mouse splenocytes and serum compared to the parents. Additionally, GEO dataset: GSE79240 was utilized to observe the differential expression of microRNAs during SLE, especially, miR-30e expression level, that further revealed that expression of miR-30e modestly enhanced in dendritic cells of SLE patients (n=5) compared to healthy controls (n=5) (Fig. S12A). To understand the relevance of miR-30e in another autoimmune disease, we reanalyzed the previously submitted GEO dataset (GSE55099) for Type 1 Diabetes Mellitus patients. The reanalysis unveils that the expression of miR-30e significantly enhanced in PBMCs of patients (n=12) compared to healthy controls (n=10) (Fig. S12B). Collectively, these results suggest that enhanced innate immune responses are strongly linked with miR-30e expression under physiological condition and it might also play pivotal role in immune-pathogenesis of SLE in both human and mouse model.

**Figure 5.**
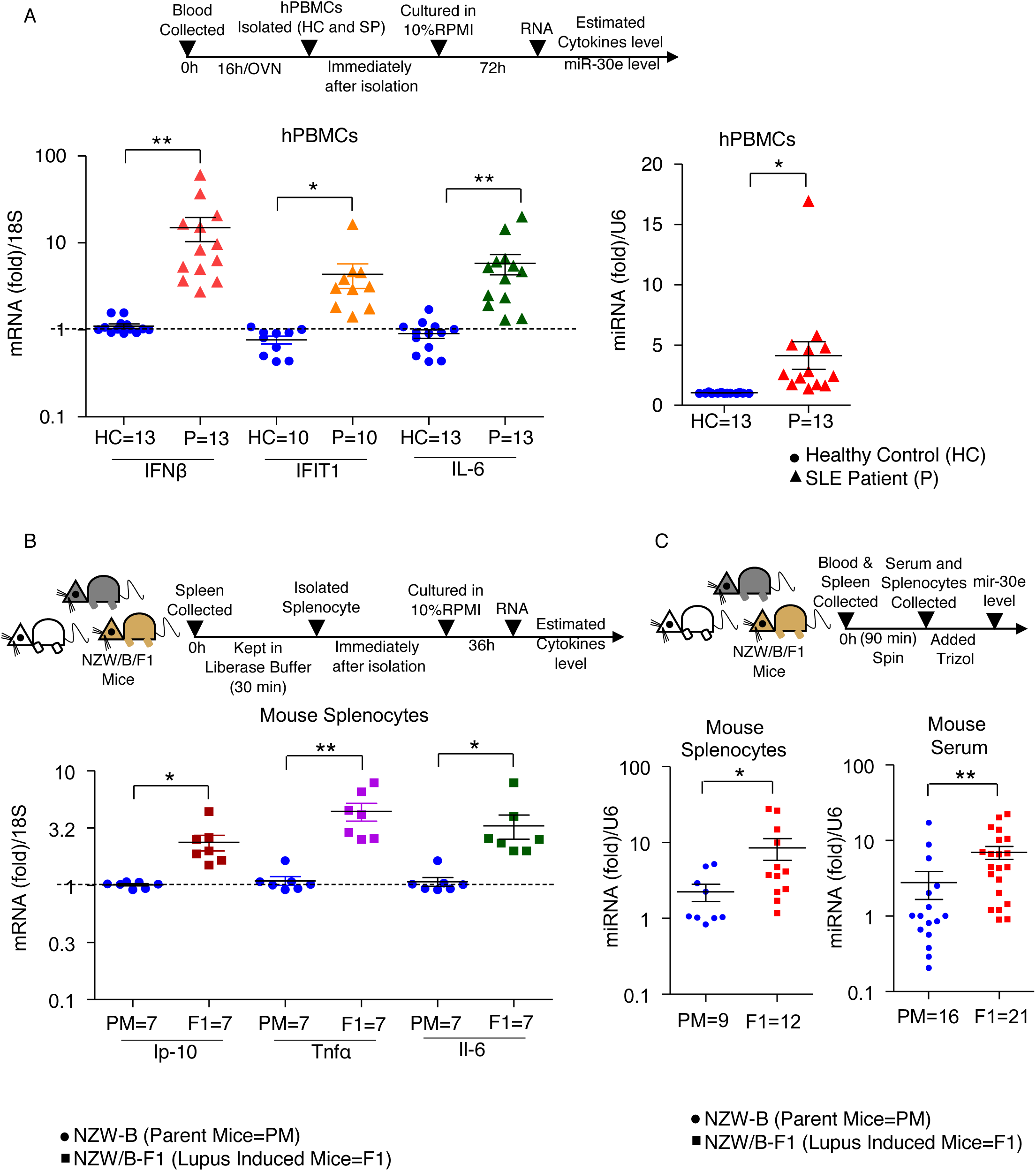
SLE patients and mouse model show enhanced innate immune responses: (A-C) Quantification of the fold changes by qRT-PCR analysis of indicated transcripts *IFNβ, IFIT1* and *IL6* (in SLE patients) and *Ip10, Tnfα* and *Il6* (in SLE mice) and miR-30e at indicated times as represented in the schematic workflow in, (A) PBMCs from SLE diagnosed patients (P) (n=13) and healthy controls (HC) (n=13), (B) Splenocytes from parent (New Zealand White and Black-NZW and NZB) (PM) (n=7) and lupus induced mice (NZW/B -F1 progeny) (F1) (n=7) and (C) Splenocytes from PM (n= 9) and F1 (n=12) and serum from PM (n=16) and F1 (n=21). Data are mean +/-SEM of triplicate samples from single experiment (A) and are representative of two independent experiments (B and C). \*\*\**P<*0.001, \*\**P<*0.01 and \**P<*0.05 by unpaired t-test and Mann-Whitney test.

### miR-30e targets negative regulators of PRR-mediated signaling pathway in SLE patients and SLE mouse model

The enhanced expression of inflammatory cytokines, type I IFNs and type I IFN-inducible genes along with elevated miR-30e in SLE patients and SLE mouse model prompted us to examine the levels of our previously identified negative regulators as shown (Fig. 3A-B). Additionally, it has been reported that several negative regulators of innate immunity play a crucial role in the development of autoimmune disease. As expected, the expression of negative regulators of PRR-mediated signaling pathways namely *TRIM38, TANK, SOCS1 and SOCS3* was significantly reduced in the PBMCs of SLE patients compared to healthy controls (Fig. 6A). To support our observation, reanalysis of previously submitted GEO dataset, GSE11909 for SLE patients (n=103) and healthy controls (n=12) in PBMCs reveal that, in SLE, the identified negative regulators of innate immune signaling pathway maybe targeted by miR-30e, were significantly reduced in patients (n=103) compared to healthy controls (n=12) (Fig. S13). To confirm our observations, splenocytes from mouse model were analyzed for identified negative regulators. Consistent with human result for the expression of negative regulators, the expression of *Atg5* and *Atg12* were significantly reduced whereas expression of *Socs1* and *Socs3* was moderately reduced in SLE mouse (F1-NZW/B mice: n=7) compared to parent mice (PM-NZW-NZB: n=7) (Fig. 6B). Collectively, these results suggest that in SLE pathogenesis, in both human and mouse, enhanced expression of miR-30e might play a crucial role, by suppressing the expression of negative regulators of innate immune signaling pathway, which in turn enhance innate immune cytokines and contribute in development or severity to the disease. Therefore, manipulation of miR-30e expression might be key for the controlling SLE pathogenesis.

**Figure 6.**
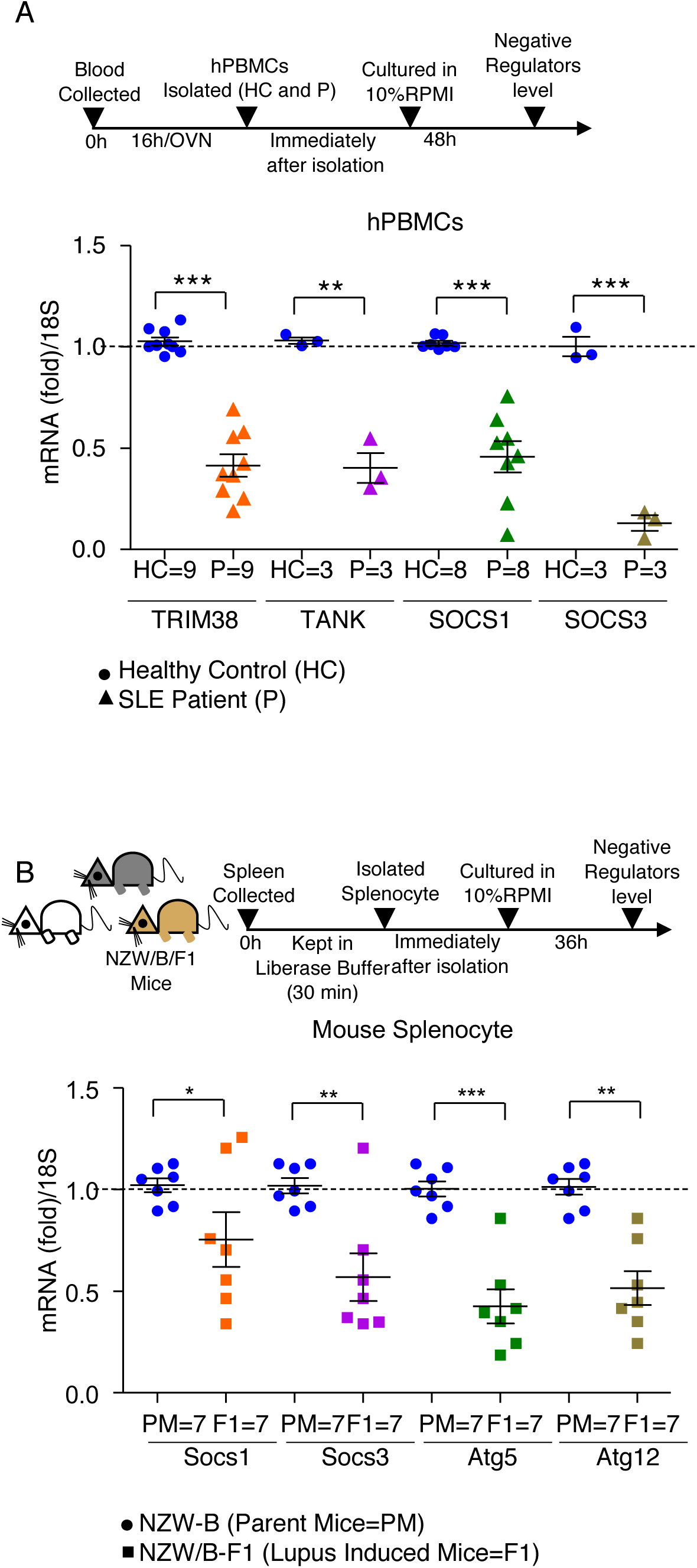
Enhanced miRNA-30e suppresses negative regulators in SLE patients and mouse model: (A-B) Schematic representation of the workflow for quantification of the fold changes by qRT-PCR analysis of indicated transcripts in (A) PBMCs of SLE patients (*TRIM38, TANK, SOCS1* and *SOCS3*) and (B) Splenocytes of SLE mice (*Socs1, Socs3, Atg5* and *Atg12*). Data are mean +/-SEM of triplicate samples from single experiment (A) and are representative of two independent experiments (B). \*\*\**P<*0.001, \*\**P<*0.01 and \**P<*0.05 by unpaired t-test.

### Prognostic and Therapeutic potential of miR-30e

To explore the prognostic and therapeutic potential of miR-30e, which modulate innate immune responses during infection and autoimmune diseases, particularly HBV infection and SLE, we tested prognostic potential of miR-30e. We validated the miR-30e expression by comparing the serum levels in pre- and post six months therapy (pegylated type I interferon) samples of HBV patients (demographic details mentioned in Table T6). Strikingly, we found significant reduction of miR-30e expression after interferon therapy (Fig. 7A), suggesting that, HBV infection triggers miR-30e over-expression in the host, to combat the which otherwise ablates upon Peg-IFN treatment, a well-established therapy that reduces the hepatitis B virus titer in patients. Additionally, GEO dataset GSE104126 supports the above finding in context to miR-30e expression level, as its reanalysis revealed, that there was significant reduction in the level of miR-30e in pegylated type I interferons treated and HBsAg-loss (reduction in viral titer) patients compared to pegylated type I interferons treated and non-HBsAg loss (no change in viral titer) patients (Fig. S14).

**Figure 7.**
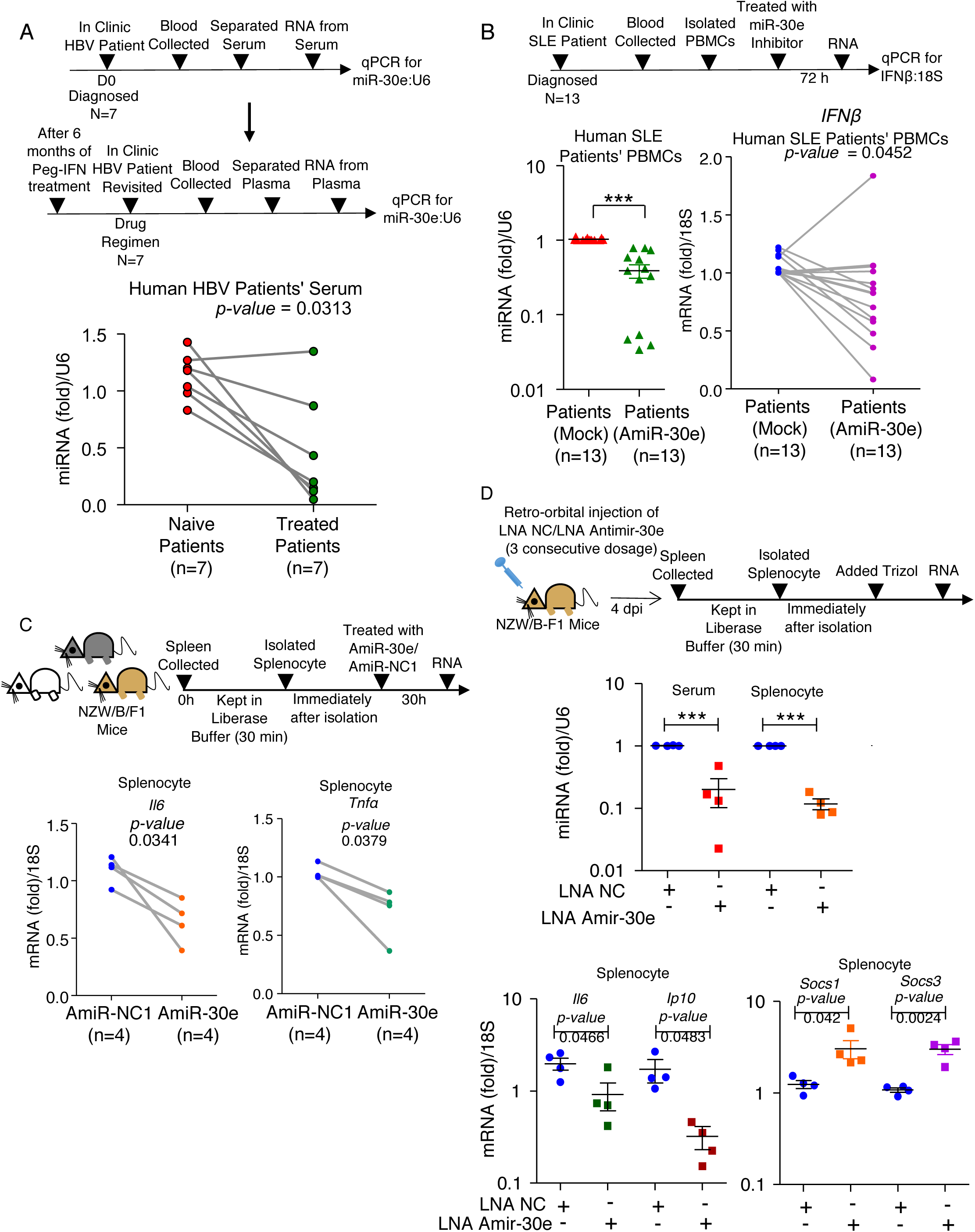
Prognostic and therapeutic potential of miR-30e: (A) Schematic representation of the workflow for quantification of the fold changes by qRT-PCR analysis of miR-30e as indicated, in the serum collected from hepatitis B (HBV) naive patients (n=7) compared to HBV treated (with pegylated IFNs) patients (n=7). 1. (B) Schematic representation of workflow for quantification of the fold changes in the relative abundances of miR-30e and *IFNβ* as indicated, in the PBMCs from SLE patients treated with/without AmiR-30e (miR-30e inhibitor). (C) Schematic representation of the *ex-vivo* experiment workflow for quantification of the fold changes in the relative abundances of *Il6* and *Tnfα* as indicated, in the splenocytes from SLE mice model (as described previously) treated with AmiR-30e (miR-30e inhibitor) and AmiR-NC1. (D) Schematic representation of the *in-vivo* experiment workflow for quantification of the fold changes of miR-30e, *Il6*, *Ip10*, *Socs1* and *Socs3* as indicated, in the splenocytes and serum from SLE mice model (as described previously); four mice distributed in each group were subjected to LNA-miR-30e-antagomir (LNA Amir-30e) and LNA-negative control antagomir (LNA NC) treatment (explained in materials and methods). Data are mean +/-SEM of triplicate samples from single experiment (A-B and D) and are representative of two independent experiments (C). All the *P-values/*\*\*\**P<*0.001 defined by paired t-test.

**Figure 8.**
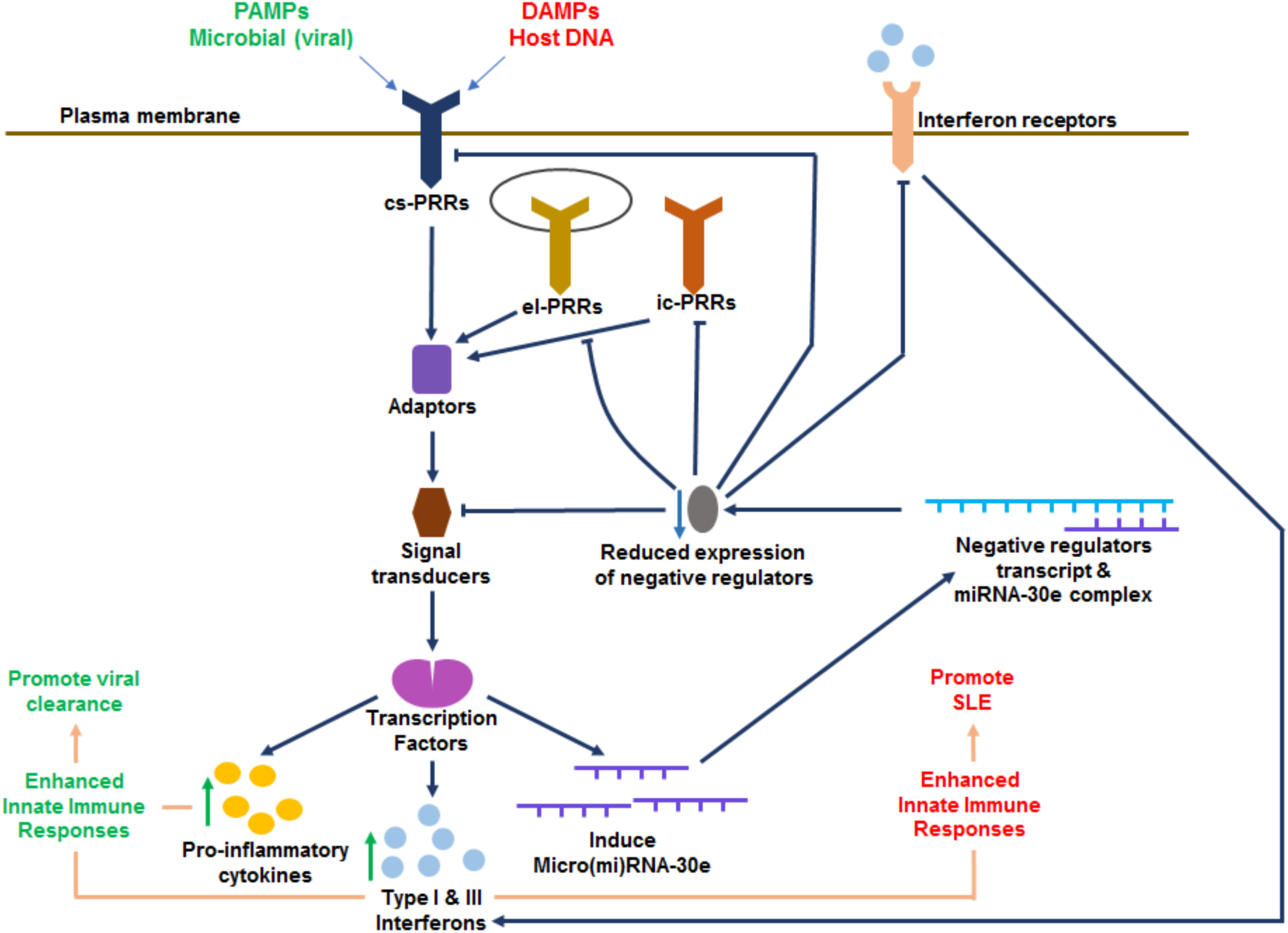
Regulation of innate immune responses by miRNA-30e during virus infection and SLE: PAMPs (green) and DAMPs (red) sensed by Pattern recognition receptors (PRRs) to activate cascade of innate immune signaling pathways to induce pro-inflammatory cytokines (yellow), type I and type III interferons (blue) and miRNA-30e (purple). miRNA-30e regulates both PAMPs and DAMPs induced immune responses by targeting the 3’-UTR of negative regulators (dark blue) of innate immune signaling pathways and reducing the expression of these negative regulators (grey). During viral infection, miR-30e is induced which reduces the cellular abundance of negative regulators to enhance innate immune responses and facilitate viral clearance. The endogenous host DNA induces miR-30e and subsequently enhances innate immune responses for the development of autoimmune disease, SLE.

Next, we investigated the therapeutic potential of miR-30e modulator and have shown the importance of AmiR-30e (miR-30e inhibitor) that sequesters the activity of endogenous miR-30e. Our results showed that SLE patients and mouse model produce high inflammatory responses in terms of innate cytokines and it is linked to the elevated miR-30e expression (Fig. 5A) and contribute in reduction of negative regulators (Fig. 6A). Therefore, we introduce AmiR-30e into SLE patient’s PBMCs. Interestingly, we found that ectopic expression of AmiR-30e sequester the expression of miR-30e in PBMCs of SLE patients and reduces the *IFNβ* expression as quantified by qRT-PCR (Fig. 7B). Next, we examined the role of AmiR-30e in SLE mouse model, NZW/B. In *ex-vivo* experiment, transfection of splenocytes derived from the F1 mice (NZW/B) with AmiR-30e significantly reduces the expression levels of *Il6 and Tnfα* compared to the control AmiR-NC1 (Fig. 7C). Furthermore, to show the stable and specific sequestering effects of microRNAs for therapeutic relevance, previously it has been published that locked nucleic acid (LNA) based chemistry to design a potent inhibitor against the microRNA has showed promising effects in *in-vivo* studies^12^. Therefore, finally, we performed *in-vivo* experiments to test the effects of LNA-anti-mir-30e-5p (LNA Amir-30e) on innate immune responses in SLE mice, two groups of mice each consists of four mice were injected with LNA Amir-30e or LNA negative control (LNA NC) through retro-orbital route thrice with one day interval to sequester the endogenous expression of mir-30e to measure mir-30e and gene expression levels. The expression of mir-30e in serum and splenocytes of SLE induced F1 (NZW/B) mice was significantly reduced compared to the LNA NC treated mice. Additionally, innate immune cytokines namely *Il6 and Ip10* significantly reduced in LNA Amir-30e treated mice compared to the LNA NC. In contrast the expression of negative regulators, *Socs1 and Socs3* significantly enhanced in LNA Amir-30e treated mice compared to the LNA NC. (Fig. 7D). Taken together these findings conclude that LNA Amir-30e found to be stable under *in-vivo* conditions and significantly inhibits the activity of mir-30e in SLE mouse model, which further reduces the inhibition/targeting of negative regulators, contributing towards controlling of SLE phenomena.

## Discussion

Innate immune responses to viral infection induces the production of pro-inflammatory cytokines and type I Interferons (IFNs) through cascade of complex signaling pathways that play critical roles in development of appropriate anti-viral immunity. In contrast, dysregulation of these signaling pathway results to inefficient clearance of microbial infection, immunopathology or autoimmune diseases^1, 3^. Therefore, the expression and activation of signaling molecules in signaling pathways are tightly regulated at transcriptional, post-transcriptional, translational and post-translational levels. The non-coding small (micro) RNAs play a pivotal role in fine tuning of protein coding genes through, stability and translation of gene transcript. Here our study identifies a novel role of miR-30e in regulation of innate immune signaling pathway during HBV infection and immuno-pathogenesis of SLE. We also demonstrated the therapeutic and prognostic potential of AmiR-30e and miR-30e in SLE and HBV infection, respectively.

The miR-30e identified through unbiased *in-silico* screening using GEO datasets obtained from cell lines, primary cells, mice or human patients challenged by different viruses. Although, other miRNAs such as miR-27a-3p and miR-181a/2-3p are also induced, however, their complementation potency towards mRNA-miRNA targets was manifolds lower than miR-30e. The miR-30e has been reported to be associated with cancer^13–15^, cardiac dysfunction^16^, kidney malfunction^17^, fatty acid deregulation, as a biomarker for SLE^18^, a dysregulated micro RNA during Zika virus infection^19^ and suppressor for Dengue virus^20^. However, its role in innate immune signaling pathway and innate immune responses during virus infection and pathogenesis of autoimmune diseases such as SLE are not clear.

Our results show that miR-30e induced manifold in the serum of (n=51) therapy naive chronic hepatitis B (CHB) patients compared to healthy control (n=24). This finding was also supported by another DNA and RNA virus such as HCMV, NDV, and SeV infection or stimulation with TLR or RLR viral PAMPs such as ssRNA and poly IC to various cell lines. Ectopic expression of miR-30e in primary cells or cell lines upon subsequent DNA or RNA virus infection significantly reduces the viral load through global enhancement of innate immune responses in terms of pro-inflammatory cytokines, type I interferons and type I interferons-inducible genes as shown by NGS data analysis. The enhanced miR-30e expression in therapy naive HBV (CHB) patients might elevate innate immune responses to combat HBV infection, however, it might be insufficient to control HBV infection. Therefore, HBV patients receiving pegylated interferon were sampled post six months therapy and were found to show significant reduction of viral load and miR-30e expression highlighting the link between HBV, miR-30e and innate immune responses. To establish the role of miR-30e under physiological condition, we selected SLE as a disease model. The SLE patients show enhanced expression of miR-30e, pro-inflammatory cytokines, type I interferons or type I interferons-inducible genes. Further to confirm our observations, we exploited SLE mouse model (NZW/ NZB F1) and obtained similar results for the expression of miR-30e that were consistent with SLE patients result suggesting the correlation of miR-30e with innate immune responses in SLE under physiological condition.

The transcriptomic analysis of our NGS data and GEO data sets using various bioinformatics tools shows that miR-30e directly targets several signal transducers and the negative regulators such as *TRIM38*^21–23^*, TANK*^24^*, ATG5, ATG12*^25, 26^*, BECN1*^27^*, SOCS1* and *SOCS3*^28^.

The microRNA targeting reduces the expression negative regulators and enhance the innate immune responses which play pivotal role in TLR, RLR, NLR, and type I interferon signaling pathways. Although miR-30e also bind with few other gene transcripts which are not negative regulator in the innate immune signaling pathway, however, the combined mean free energy for these transcripts are lower than threshold binding energy necessary for significant change in the expression of transcripts, for subsequently affecting the outcome of signaling pathway. Notably, the expression of negative regulators such as *TRIM38, TANK, SOCS1* and *SOCS3* in SLE patients are significantly reduced. Moreover, SLE mice also show similar results for the expression of *Socs1, Socs3, Atg5* and *Atg12*. Collectively, human and mouse results illustrate that miR-30e targets negative regulators to elevate innate immune responses and dysregulation of miR-30e expression may be one of factor for the establishment of SLE or other autoimmune diseases.

Previously, it has been shown that SLE patients has reduced ability to degrade DNA and cellular chromatin^29^. Therefore, PBMCs from healthy donors were stimulated with partially degraded self-DNA and these cells showed enhanced miR-30e and innate immune responses suggesting the pathogenic role of miR-30e and suggested a link of self-DNA with the pathogenesis of SLE. In contrast, sequestering endogenous miR-30e in PBMCs of SLE patients by introducing antagomir/inhibitor significantly reduced the levels of miR-30e and innate immune responses in terms of *IFNβ* production. Additionally, introduction of locked nucleic acid (a stable form of antagomir) through retro-orbital route into SLE mice model significantly reduced mir-30e and innate immune genes expression in splenocytes whereas expression of negative regulators was enhanced, demonstrating the ability of mir-30e antagomir to reduce innate immune responses under physiological condition. Finally, our study identified miR-30e as a post-transcriptional regulator of negative regulation of PRR-mediated innate immune signaling pathways and its diagnostic and prognostic potential in HBV infection. Additionally, our study demonstrated the therapeutic implications of miR-30e antagomir/inhibitor to immunologically complex autoimmune disease, SLE or possibly other autoimmune diseases.

## Materials and Methods

### Ethical Statement

Experiments were performed after approval from the Institutional Ethical Committee (IEC)-Indian Institute of Science Education and Research (IISER) Bhopal: (IISERB/IEC/Certificate/20l6-IV/03), Institute Biosafety Committee (IBSC) - IISER-Bhopal: (IBSC/IISERB/2018/Meeting II/08), Bhopal Memorial Hospital & Research Centre Institutional Ethical Committee (IEC), BMHRC Research Projects/Clinical Studies (IRB/18/Research/10), Institutional Animal Ethical Committee (IAEC)-Small and Experimental Animal Facility National Institute of Immunology (NII): (IAEC#469/18) and Institutional Ethical Committee (IEC)-Sanjay Gandhi Post Graduate Institute of Medical Sciences (SGPGIMS): (2016-138-EMO-93).

### Human blood samples

Blood samples from both healthy individuals and patients (HBV and SLE) were collected according to the ethics protocol in the respective hospitals by health professionals. Written informed consent was obtained from all patients and healthy participants before inclusion into the study at Bhopal Memorial Hospital & Research Center (BMHRC: for HBV samples: n = 51 Vs control samples: n = 24), Sanjay Gandhi Post Graduate Institute of Medical Sciences (SGPGIMS: for SLE samples: n = 13 Vs control samples: n = 13) and Institutional Ethical Committee (IEC)-Indian Institute of Science Education and Research (IISER) Bhopal. Reagents used for transfection/electroporation were Lipofectamine 2000, miRNA mimics, miRNA inhibitors, controls mimic/inhibitors and for RNA isolation were Trizol/Trizol-LS. All reagents used were procured from Ambion/Invitogen.

### Mice

Systemic Lupus Erythematous (SLE) mouse model were procured from the Jackson Laboratory, USA and further breeding was done in approved pathogen-free, small animal facility of National Institute of Immunology (NII). SLE mouse strains were used as follows: New Zealand White (NZW) and New Zealand Black (NZB) non-lupus bearing parent mice crossed to generate NZW/B F1 progenies, lupus induced mice. All the mice used in the study were from 6 to 12 weeks without any genders bias. Dendritic cell enriched low denisty fraction of splenocytes were prepared as described earlier^40, 41^. Spleens were harvested and single cell suspension of splenocytes were cultured in RPMI based media containing either 50nM of miR-30e antagomir and miR-NC1 antagomir (transfected through RNAimax transfecting reagent) and plated in 24-well plate (3 × 10^6^ cells/well) maintained at 37°C+5% CO_2_. For the *in-vivo* experiments, four-6 to 12 weeks old NZW/B F1 mice were randomly assigned into two groups. The mice in each group received four consecutives intravenous (retro-orbital) injections of either LNA antimiR-30e or LNA negative control (scrambled) compounds, formulated in TE Buffer (10mM Tris (pH: 7.5), 0.01mM EDTA) as per 25 mg LNA/kg mouse body weight on consecutive days as described previously. The mice were sacrificed within 24 h after the last dose. At selected time points, cells were harvested and relative abundances of miR-30e and other transcripts were quantified using qRT-PCR.

### Cell lines, virus infections, transfection and reagents

A549 human alveolar basal epithelial cells (Cell Repository, NCCS, India), HEK293T/HEK293 human embryonic kidney cells (ATCC CRL-3216), Raw 264.7 (Cell Repository, NCCS, India), HeLa cervical cancer cells (Cell Repository, NCCS, India), HepG2-NTCP cells (from Dr. Takaji Wakita National Institute of Infectious Diseases Tokyo, Japan), HepG2 hepatoblastoma cells (from Dr. Nirupma Trehanpati’s Laboratory, Institute of Liver & Biliary Sciences ILBS, New Delhi, India), HepG2215, HBV stably expressing hepatoblastoma cells (from Dr. Senthil Kumar Venugopal’s Lab, South Asian University, New Delhi, India) and HFFs (from Professor. Wade Gibson’s Lab, Johns Hopkins, School of Medicine) were cultured in Dulbecco’s modified Eagle’s medium (DMEM) supplemented with 10% fetal bovine serum (FBS) and 1% Antibiotic-Antimycotic solution. HepG2-NTCP cells were infected with HBV in Professor Akinori Takaoka’s Laoboratoy as per the standard protocol. Human PBMCs were isolated from whole blood and NDV LaSota viral stocks were accumulated as described previously^7^. NDV-GFP was a kind gift from Professor Peter Palese, Icahn School of Medicine at Mount Sinai. Sendai-RFP (SeV-RFP) were borrowed from Dr. Sunil Raghav, ILS, Bhubaneshwar, India, A single virus stock was used for all experiments. The cells were infected in serum-free DMEM with the NDV and SeV viruses at the MOIs indicated in the figure legends. After 60 min, the cells were washed with phosphate-buffered saline (PBS) and then were resuspended in DMEM, 1% FBS. HFFs were grown to full confluence and infected with GFP tagged HCMV at the MOIs indicated in the figure legends. HBV-positive sera were used to infect HepG2 cells as previously described^42–46^. HepG2 cells were made permissive to HBV virus infection by adding 3% PEG (polyethylglycol) and 0.5% DMSO (dimethylsulphoxide). The cells were infected in 1% FBS containing DMEM, 3% PEG and 0.5% DMSO with the HBV-positive sera. After 6-7 hours, the cells were washed with phosphate-buffered saline (PBS) and then were resuspended in DMEM, 10% FBS, 3% PEG and 0.5% DMSO. For electroporation of human PBMCs, 1 × 10^6^ cells were suspended in Opti-MEM (Invitrogen) containing 50 nM mirVana miRNA mimics (Ambion). The cells were pulsed twice with 1000 V for 0.5 ms with a pulse interval of 5 s with the Gene Pulser Xcell electroporation system. The cells were then transferred to RPMI supplemented with 10% FBS^7^. Transfection of cells with miRNA mimics, inhibitors and control mimics/inhibitors and/or plasmids was performed with Lipofectamine 2000 or 3000 (Invitrogen) according to the manufacturer’s protocol. Poly IC (Invivogen) was mixed with Lipofectamine2000 before being used to transfect cells. ssRNA (Invitrogen) were used to stimulate the cells as mentioned in the figure legends. DMEM, FBS, Opti-MEM, RPMI, and Lipofectamine 2000/3000 were purchased from Invitrogen. The miR-30e mimic (miR-30e) (Invitrogen) or a nonspecific miRNA negative control (miR-NC1) was used according to the manufacturer’s instructions (Applied Biosystems). The miR-30e inhibitor (AmiR-30e) (Invitrogen) was used to inhibit miR-30e expression in transfected cells. The cDNA encoding the 3′-UTR of negative regulators was retrieved from the UCSC gene sorter and was sub-cloned into the pMIR-REPORT luciferase vector. A total of 2.0 kb of sequence upstream of the miR-30e gene was retrieved from the UCSC genome browser. This sequence was amplified by PCR from genomic DNA and was subcloned into the pGL3 basic vector between the KpnI and HindIII sites. Plasmids containing Firefly Luciferase gene under *IFNβ, ISRE* and *NFκB* promoters, were obtained from Professor Shizuo Akira’s (Osaka University, Japan), *rhIFNβ* (bei resources) and *rhTNFα* (R&D Systems). All sh-clones, were obtained from the whole RNAi human library for shRNA mediating silencing (Sigma, Aldrich) maintained at IISER, Bhopal, India. *In-silico* analysis for miRNA target gene prediction was done as previously described^7^.

### Quantitative real-time reverse transcription PCR

Total RNA was extracted with the Trizol reagent (Ambion/Invitrogen) and used to synthesize cDNA with the iScript cDNA Synthesis Kit (BioRad, Hercules, CA, USA) according to the manufacturer’s protocol. Gene expression was measured by quantitative real-time PCR using gene-specific primers and SYBR Green (Biorad, Hercules, CA, USA). For quantification of the abundances of miR-30e, real-time PCR analysis was performed with the TaqMan Universal PCR Master Mix (Applied Biosystems) and the miR-30e-5p specific TaqMan miRNA assays. The Taqman U6 assay was used as a reference control. Real time quantification was done using StepOne Plus Real time PCR Systems by Applied BioSystems (Foster City, CA, USA). Primers used for qRT-PCR were listed in Supplementary Table T7.

### Luciferase Reporter assays

HEK 293T and HeLa cells (5 × 10^4^) were seeded into a 24-well plate and transiently transfected with 25 nM of mimics, 50 ng of the transfection control pRL-TK plasmid (*Renilla* luciferase containing plasmid) and 200 ng of the luciferase reporter plasmid (*Firefly* luciferase containing plasmid) together with/without 300 ng of the various expression plasmids or an empty plasmid as a control according to the respective experiments. The cells were lysed at 24 to 36 hours after transfection and/or infection or stimulations, and finally the luciferase activity in total cell lysates was measured with Glomax (Promega, Madison, WI, USA).

### Enzyme-linked immunosorbent assay (ELISA)

A549 and HeLa cells were transiently transfected with miR-30e and miR-NC1 and then were infected NDV virus. The culture media were harvested 36 to 40 hours after infection and were analyzed by specific ELISA kits (Becton Dickinson) according to the manufacturer’s instructions to determine the amounts of *IP10* and *IL6* that were secreted by the cells.

### RNA immunoprecipitations

RNA immunoprecipitations were performed as described previously^47, 48^. The pIRESneo-Flag/HA Ago2 plasmid was a gift from Professor T. Tuschl (Addgene plasmid #10822). Briefly, HEK 293T cells transfected with miRNA and infected with NDV were lysed in 0.5% NP-40, 150 mM KCl, 25 mM tris-glycine (pH 7.5) and incubated with M2 Flag affinity beads (Sigma) overnight. The lysate was then washed with 300 mM NaCl, 50 mM tris-glycine (pH 7.5), 5 mM MgCl_2_, and 0.05% NP-40. The extraction of RNA from the immunoprecipitated RNPs was performed with the Trizol reagent (Ambion, Invitrogen) according to the manufacturer’s protocol.

### Fluorescence-activated cell sorting (FACS) Cytometry Analysis

A549 and HeLa cells were grown to 70-80% confluence, then treated with mimics and negative control reagents and finally infected with SeV-RFP and NDV-GFP. After 24 hours of infection, cells were trypsinized, harvested and then washed with PBS thrice and finally resuspended in PBS for FACS analysis as described in figure legends. Human PBMCs were treated with DNA and or camptothecin (0.3μM) to estimate apoptosis levels. At desired time points, cells were analyzed by staining with FITC-labeled Annexin V and propidium iodide (Becton Dickinson, USA) as per manufacturer’s instructions and stained cells were analyzed using a FACS Aria III (Becton Dickinson) and data were analyzed by using FlowJo software (FlowJo, Ashland, OR, USA).

### Immunoblotting analysis

After cells were transfected with miRNA mimic and controls then after infected with NDV and/or NDV-GFP (as indicated in figures), lysates were collected and subjected to western blotting analysis as previously described^7, 49^. Cells were harvested after 36 hours of infection with standard ice-cold cell lysis buffer supplemented with 1 X protease inhibitor cocktail (obtained from Sigma, Aldrich). Immunoblotting were done as previously described^49^. Immunoblotted nitrocellulose membrane was imaged with LI-COR system. Anti–GFP antibody was obtained from Sigma-Aldrich, anti-TRIM38 (from ImmunoTag,), anti-TANK, SOCS3, BECN1 (from Cloud-Clone Corporation), anti-ATG12 (from Cell Signaling Technology), anti-SOCS1 (from Santa Cruz) and anti *γ*-Tubulin (from Sigma, Aldrich). IR dye labeled anti-Rabbit and anti-Mouse IgG (secondary antibody), were purchased from LI-COR.

### Microscopy

HeLa cells were transfected with miRNA mimic and infected with NDV-GFP were fixed with 4% PFA for 15 min at room temperature; permeabilized with 0.05% Triton X-100 in 1 x PBS for 10 min at room temperature; blocked with bovine serum albumin (5 mg/ml) in PBS, 0.04% Tween 20 for 30 min and incubated for 1 hour with the relevant primary antibodies diluted in blocking buffer. The cells were then washed three times with PBS and incubated for 1 hour with the appropriate secondary antibodies at room temperature. Nuclei were stained with DAPI, and the cells were then analyzed with an LSM 780 confocal laser microscope (Carl Zeiss). The images were analyzed using ImageJ processing software. HCMV infection (GFP fluorescence) in HFFs miRNA mimic and control mimic transfected cells was visualized with Inverted microscope Vert.A1 (AXIO) by Zeiss.

### RNA-Sequencing data analysis

Trizol reagent (Ambion, Invitrogen) was used to isolate total RNA that was processed to prepare cDNA libraries using TruSeq technology according to the manufacturer’s instructions protocol (Illumina, San Diego, CA). Libraries were sequenced using Illumina NovaSeq 6000, with a read length of 101 bp, by Bencos Research Solutions Pvt. Ltd., Bangalore, India. FastQC (0.11.5) was used to access the read quality of the raw data. Trimmomatic was used to remove Illumina adaptors and sliding-window approach was used for the quality filtering of reads. Approximately 20 million cleaned pair-end sequencing reads from each sample were uploaded to the Galaxy web platform and were analyzed at https://usegalaxy.org. HISAT2 was used to map the reads with the reference human genome (hg38). StringTie was used to assemble the aligned RNA-Seq reads into transcripts and estimate the abundance of the assembled transcripts. DESeq2 was used for differential expression analysis of genes between groups^30^. Various R packages were used to visualize the expression and differential expression outcomes. Gene ontology (GO) analysis was done using the web-based Gene Set Analysis toolkit, and analysis of upregulated KEGG pathways was done using Enrichr. Cluster 3.0 and TreeView 1.1.6 were used for making heat maps. All the addressed analysis were demonstrated as described previously^50^.

### Statistical analysis

All experiments were carried out along with the appropriate controls, indicated as untreated/untransfected cells (Ctrl) or transfected with the transfection reagent alone (Mock). Experiments were performed in duplicates or triplicates for at least two or three times independently. GraphPad Prism 5.0 (GraphPad Software, La Jolla, CA, USA) was used for statistical analysis. The differences between two groups were compared by using an unpaired two-tailed Student’s t-test and/or Mann Whitney test additionally the paired data was analyzed using paired t-test and/or Wilcoxon sign rank test. While the differences between three groups or more were compared by using analysis of variance (ANOVA)w ith Tukey test. Differences were considered to be statistically significant when *P < 0.05*. Statistical significance in the figures is indicated as follows: *\*\*\*P < 0.001, **P < 0.01, *P < 0.05; ns*, not significant.

## Acknowledgments

We acknowledge Professor Akinori Takaoka for valuable discussions. We thank Dr. Takaji Wakita for providing the HepG2-NTCP cell lines. We thank Dr. Nirupma Trehanpati and Dr. Senthil Kumar Venugopal for providing the HepG2 and HepG2215 cell lines. We thank Dr. Sunil Raghav for providing Sendai-RFP, Professor. Peter Palese for providing NDV-GFP and Professor. Wade Gibson for providing HCMV-GFP and HFFs. We thank Professor. T. Tuschl for providing the Ago2-Flag construct through Addgene and BEI Resources for providing human rIFN*β*. We are grateful to Indian Institute of Science Education and Research (IISER) Bhopal for providing the Central Instrumentation Facility. We also thank all members of the laboratory for helpful discussions. Finally, we are eternally grateful to all the patients and healthy donors for proving their blood samples for the study.

## Author contributions

R.M. and H.K. conceptualized the study and designed the experiments; R.M. performed the experiments; S.B. and D.K. performed the mutation experiments; R.M., A.K. and H.K. analyzed the data; R.M., P.G. and H.K. designed the HBV experiments; P.G. provided the HBV patients samples; R.M., K.N. and P.G. performed the HBV experiments; R.M., A.A., P.T. and H.K. designed the SLE experiments; A.A. provided the SLE patients samples and executed SLE patient samples experiments; R.M. and A.K. performed the SLE *in-vitro* experiments; B.S.R., R.M. and P.T. performed the SLE mice related experiments; A.C. helped in procuring critical reagents; R.M. and H.K. wrote the manuscript; and H.K. supervised the entire project.

## Data and materials availability

The NGS (RNA-Sequencing) data for expression profiling reported in this paper have been deposited in the GenBank database (accession no. GSE130005).

## Supplementary Figure Legends

**Figure S1.**
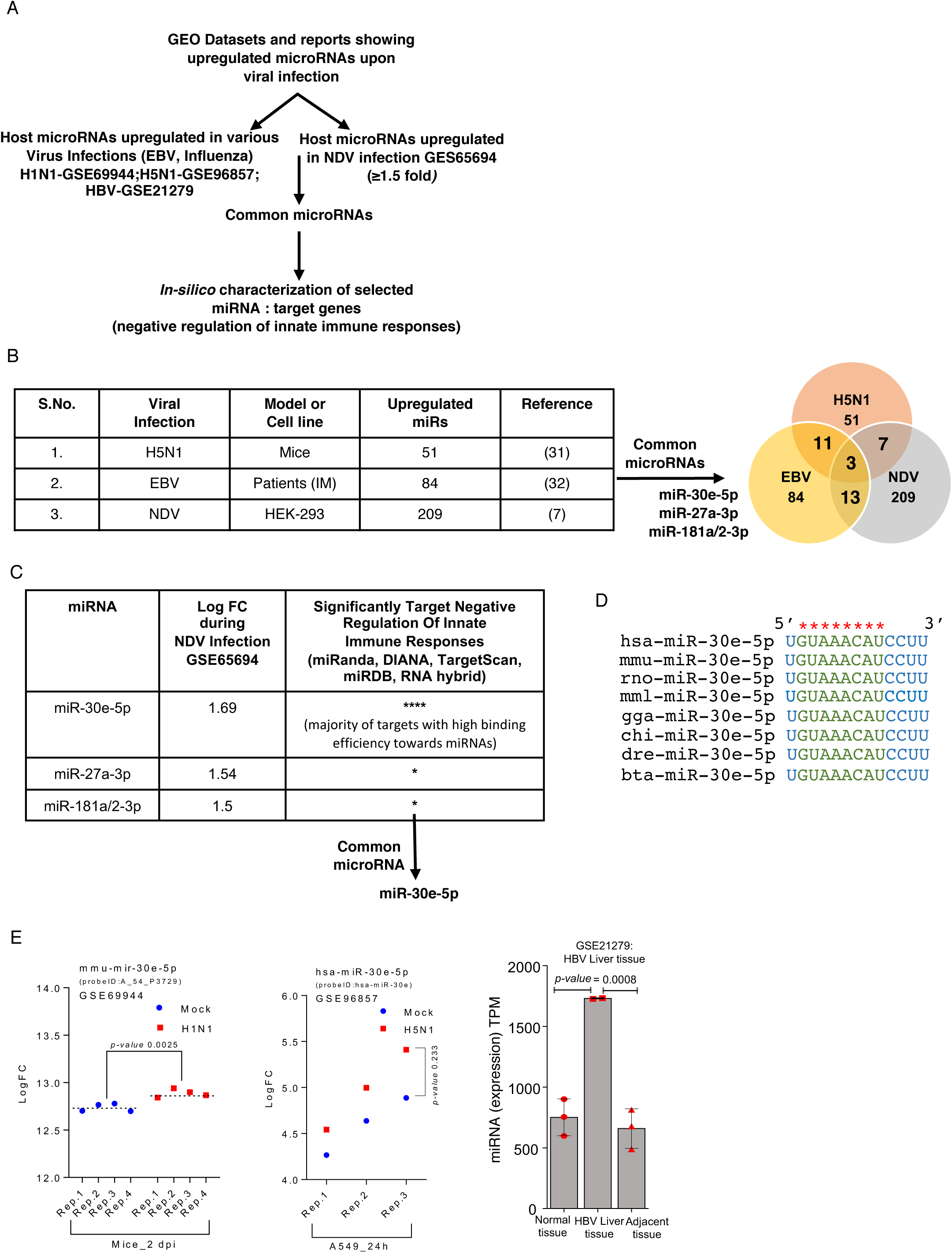
miRNA-30e induced during viral infection: (A) Schematic representation for the selection and screening pipeline of common miRNAs during viral infections. (B) Table and Venn diagram represent miR-30e, miR-27a and miR-181a as commonly upregulated miRNAs during indicated infections. Abundance of miR-30e-5p as log fold change (Log FC) and transcripts per million (TPM) in indicated infections, (C) Log FC (fold change) of the selected miRNAs and efficiency (represented by [*] asterisk) by which they target the negative regulation of innate immune signaling pathways as per indicated GEO dataset (GSE65694) and algorithms. (D) miR-30e is conserved among the wide range of species (green); has, Homo sapiens (Human); mmu, Mus musculus (Mouse); rno, Rattus norvegicus (Norway Rat); mml, Macaca mulatta (Rhesus monkey); gga, Gallus gallus (chicken); chi, Capra hircus (Goat); dre, Danio rerio (Zebrafish) and bta, Bos Taurus (Cattle).

**Figure S2.**
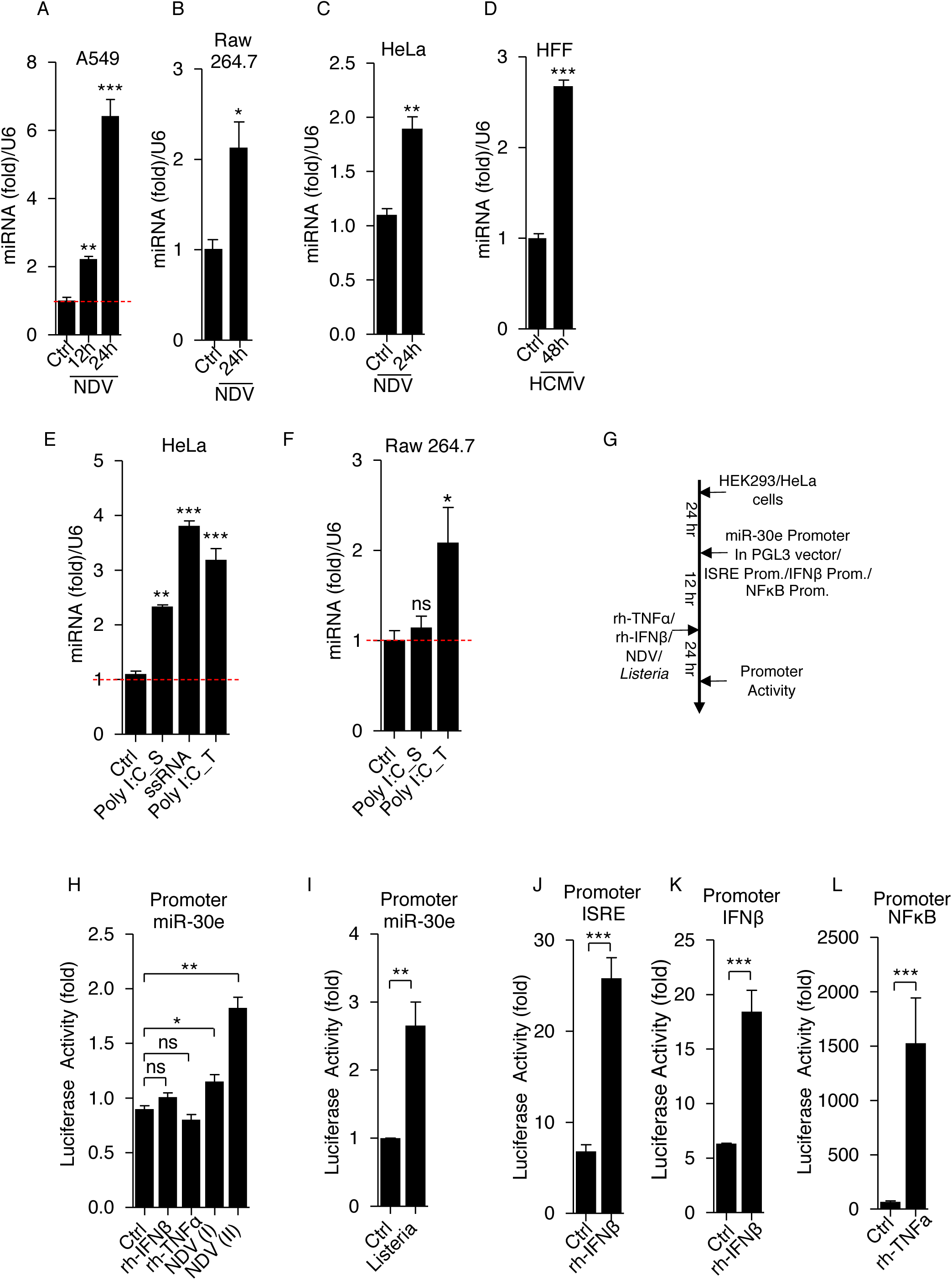
Induction of miRNA-30e in different cells by DNA, RNA virus and viral PAMPs: (A-E) Quantification of the fold changes by qRT-PCR in the abundances of miR-30e (at the indicated times and cells) after indicated viral infections and viral PAMPs treatment (A and B) NDV (MOI 5) in A549 and Raw 264.7 cells respectively, (C) HCMV-GFP (MOI 5) in HFF. (D and E) indicated synthetic PAMPs [poly IC (10μg/ml) {stimulation (S) and transfection (T)}; ssRNA (2μg/ml)] in (D) HeLa and (E) Raw 264.7 cells. (F) Schematic representation of workflow for quantification of miR-30e promoter activity and ISRE/IFN*β*/NF*κ*B promoter activity by luciferase assay as indicated in (G, I-K) HEK293 cells and (H) HeLa cells. Data are mean +/-SEM of triplicate samples from single experiment and are representative of three (A-C, I-M) two (D-G) independent experiments. \*\*\**P<*0.001, \*\**P<*0.01 and \**P<*0.05 by one-way ANOVA Tukey test and unpaired t-test.

**Figure S3.**
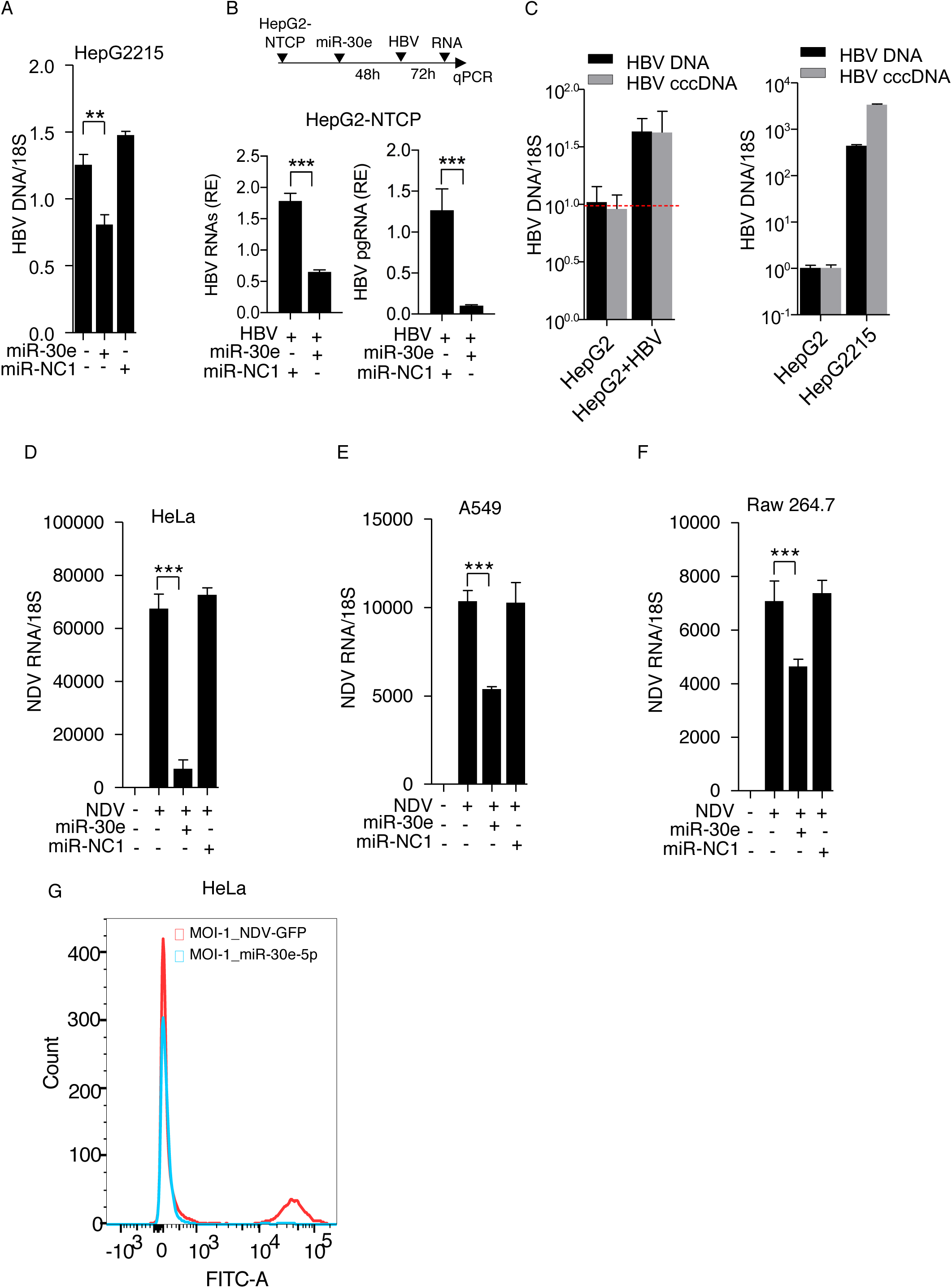
miR-30e inhibits viral replication: (A-F) Quantification of the fold changes in the relative abundances of viral transcripts measured by qRT-PCR after treatment or infection in indicated cells with (A) HepG2215 cells were transfected with/without miR-30e as described previously, (B) HepG2-NTCP cells transfected with miR-30e and miR-NC1 to quantify relative expression (RE) of HBV viral transcripts (HBV RNA and pgRNA), (C) HBV patient serum in HepG2 compared with HepG2215 cells as indicated. (D-F) NDV (MOI 5) in HeLa, A549 and Raw264.7 cells respectively transfected with miR-NC1 or miR-30e prior to infection as described previously. (G) Quantification of NDV viral signals detected by flow cytometery in HeLa cells mock transfected or transfected with miR-30e for 24 hours then subjected to NDV (GFP tagged) infection (MOI 5) for 24 hours. HBV DNA, HBV cccDNA and HBV copy number represent different primers set used to measure HBV viral transcripts. Data are mean +/-SEM of triplicate samples from single experiment and are representative of three (D-F) two (A, B,C and G) independent experiments. \*\*\**P<*0.001 and \*\**P<*0.01 by one-way ANOVA Tukey test and unpaired t-test.

**Figure S4.**
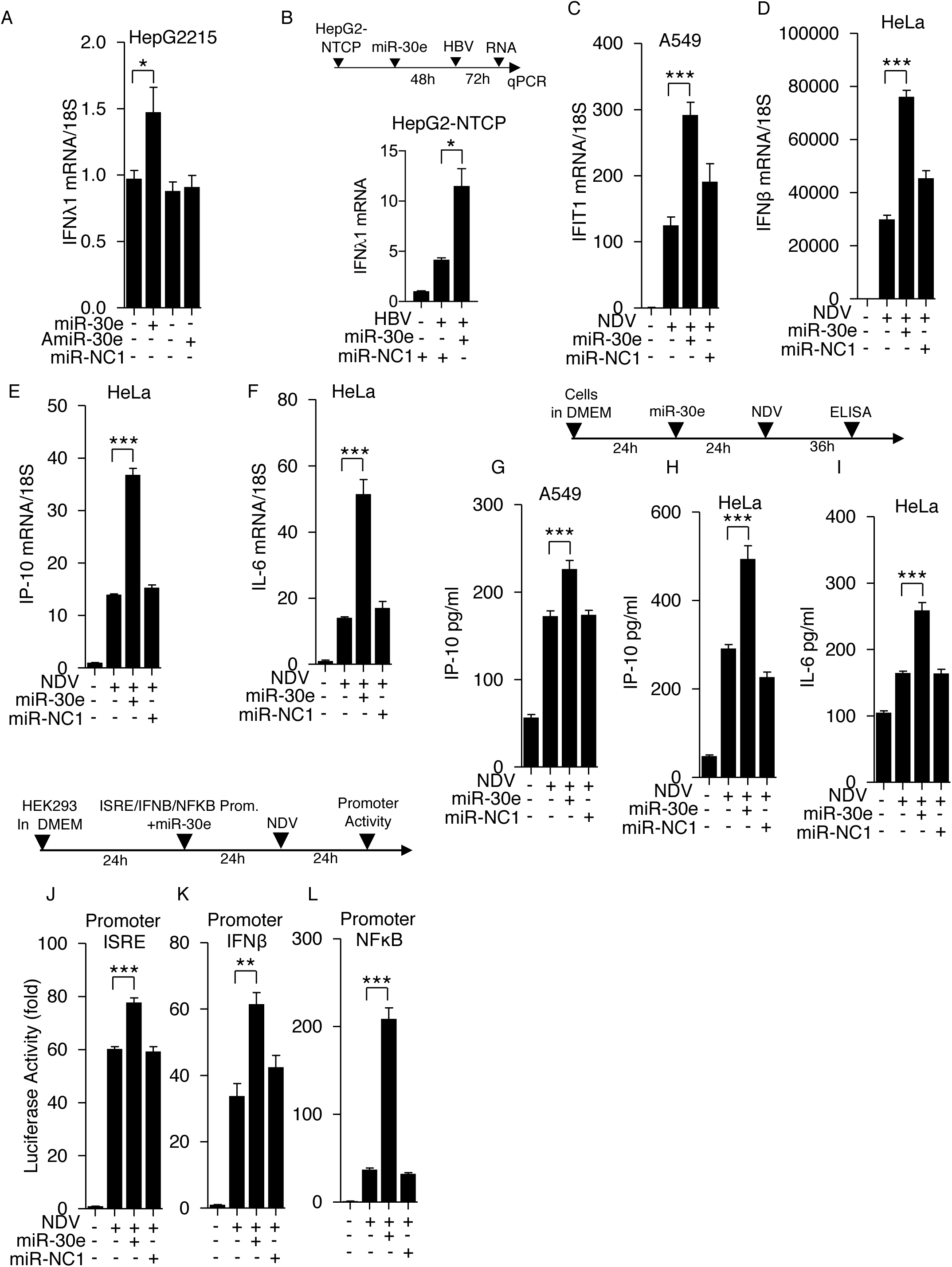
miR-30e enhances innate immune responses during viral infection. (A-F) Quantification of the fold changes in the relative abundances of indicated genes and cells measured by qRT-PCR after viral infection and transfection with/without miR-30e as described previously. (G-I) A549 and HeLa cells were mock transfected, transfected with miR-30e or miR-NC1 mimics and then infected with NDV (MOI 5), 24 hours after infection, the amounts of *IP10* and *IL6* protein secreted into the cell culture supernatant were measured by enzyme-linked immunosorbent assay (ELISA). (J-L) Schematic representation of workflow for quantification of ISRE/IFN*β*/NF*κ*B promoter activity by luciferase assay as indicated in (I-K) HEK293 cells. Data are mean +/-SEM of triplicate samples from single experiment and are representative of two (A-B) and three (C-L) independent experiments. \*\*\**P<*0.001, \*\**P<*0.01 and \**P<*0.05 by one-way ANOVA Tukey test.

**Figure S5.**
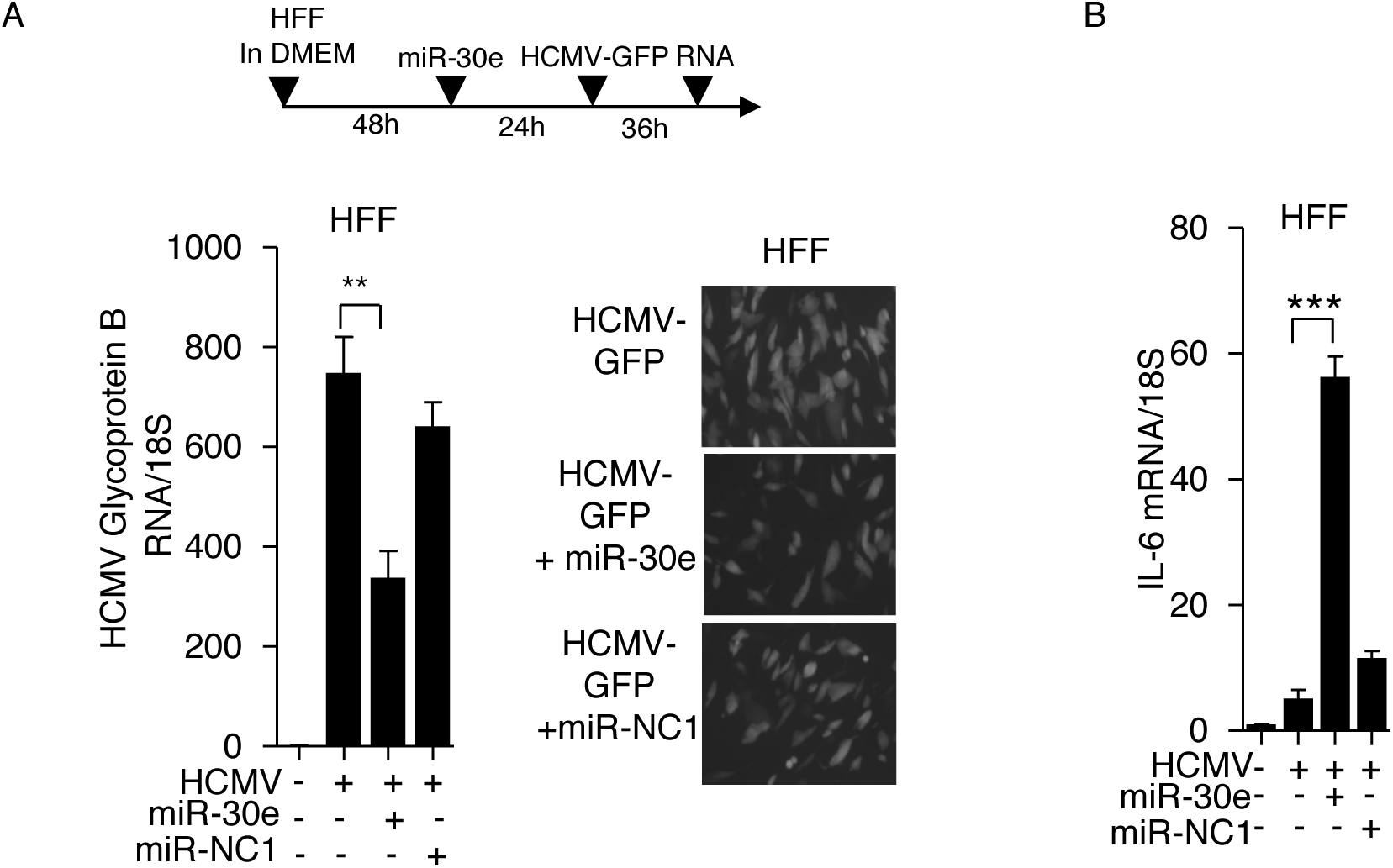
miR-30e inhibits DNA virus replication: HFF (Human foreskin fibroblast) cells were cultured in DMEM and transfected with miR-30e or miR-NC1 mimics then infected with HCMV-GFP virus (MOI = 5) and RNA were isolated to quantify the (A) HCMV transcript (Glycoprotein B) by using qRT-PCR and GFP signals (of HCMV-GFP tagged virus) by microscopy, (B) IL6 transcript in indicated transfected groups and compared with control. Data are mean +/-SEM of triplicate samples from single experiment and are representative of two independent experiments. ***P<0.001 and **P<0.01 by one-way ANOVA Tukey test.

**Figure S6.**
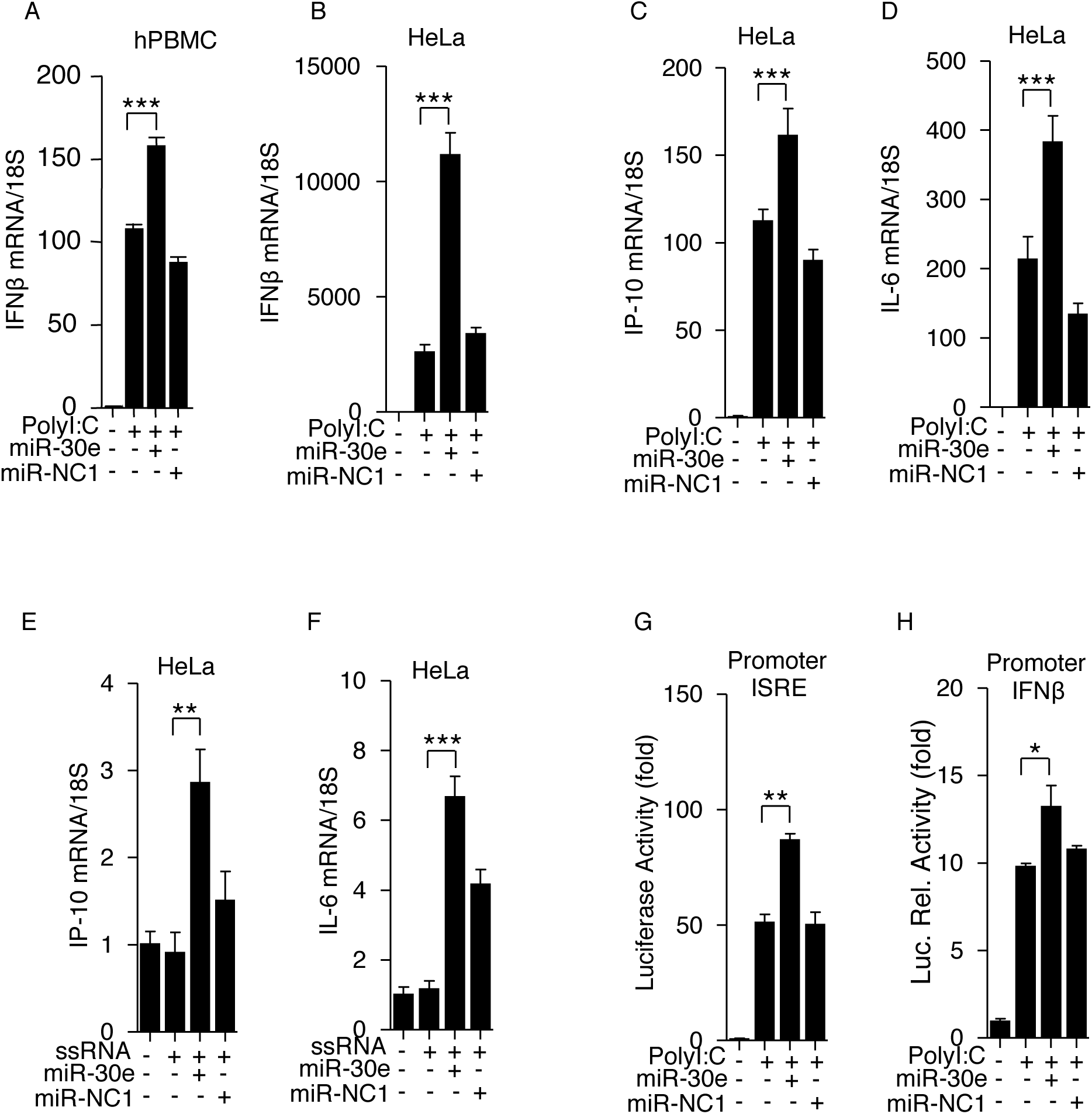
miR-30e enhances innate immune responses upon viral PAMPs stimulation: (A-F) Quantification of indicated transcripts in indicated cells (measured by qRT-pCR) transfected with mir-30e or miR-NC1 for 24 hours and then stimulated with (A-D) poly IC (10μg/ml) and (E-F) ssRNA (2μg/ml) treatment respectively. (G, H) HEK 293T cells transfected with miR-30e (25 nM) or miR-NC1 (25 nM) mimics, pRL-TK (50 ng) and indicated luciferase reporters for ISRE/IFN*β* (200 ng) then stimulated with poly IC for 24 hours. Cells were then lysed to analyze the promoter activity by luciferase assay. Data are mean +/-SEM of triplicate samples from single experiment and are representative of three independent experiments. \*\*\**P<*0.001, \*\**P<*0.01 and \**P<*0.05 by one-way ANOVA Tukey test.

**Figure S7.**
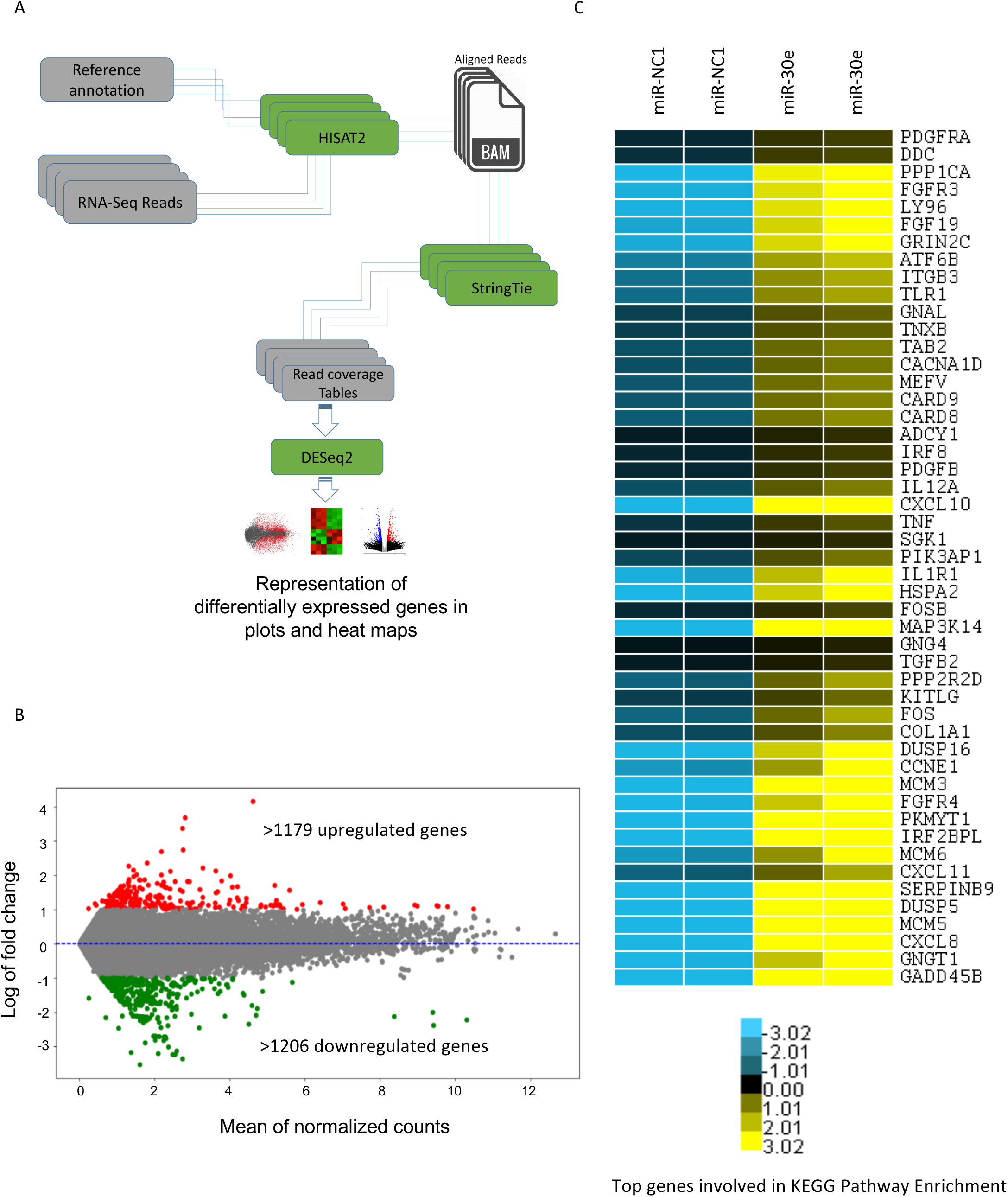
miR-30e differentially expressed genes during NDV infection: (A) Workflow for the analysis of RNA-Sequencing data from raw sequencing reads to expression profiles of differentially expressed genes represented through different plots and heat maps. (B) MA (M=log ratio and A=mean average) plot for differential expression of genes, indicating upregulated (in red) and downregulated (in green) genes. (C) Heat map representing relative abundance of genes involved in top enriched KEGG pathways in figure 2E.

**Figure S8.**
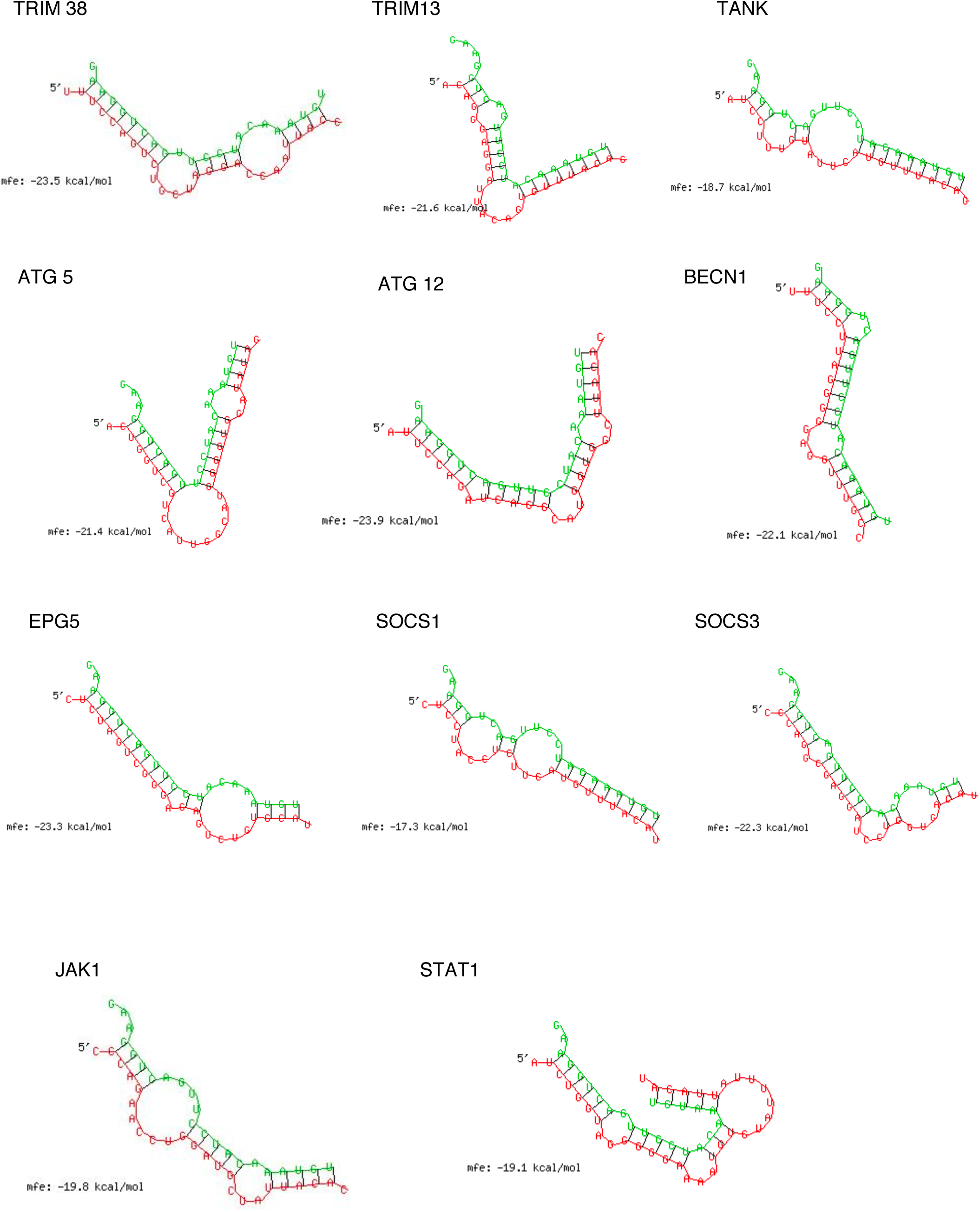
Minimum free energy (mfe) for binding efficiency of miR-30e to negative regulators: Representative of minimum free energy diagrams for negative regulators (*TRIM38, TRIM13, TANK, ATG5, ATG12, BECN1, EPG5, SOCS1* and *SOCS3*) and positive regulators (*JAK1* and *STAT1*) targeted by miR-30e.

**Figure S9.**
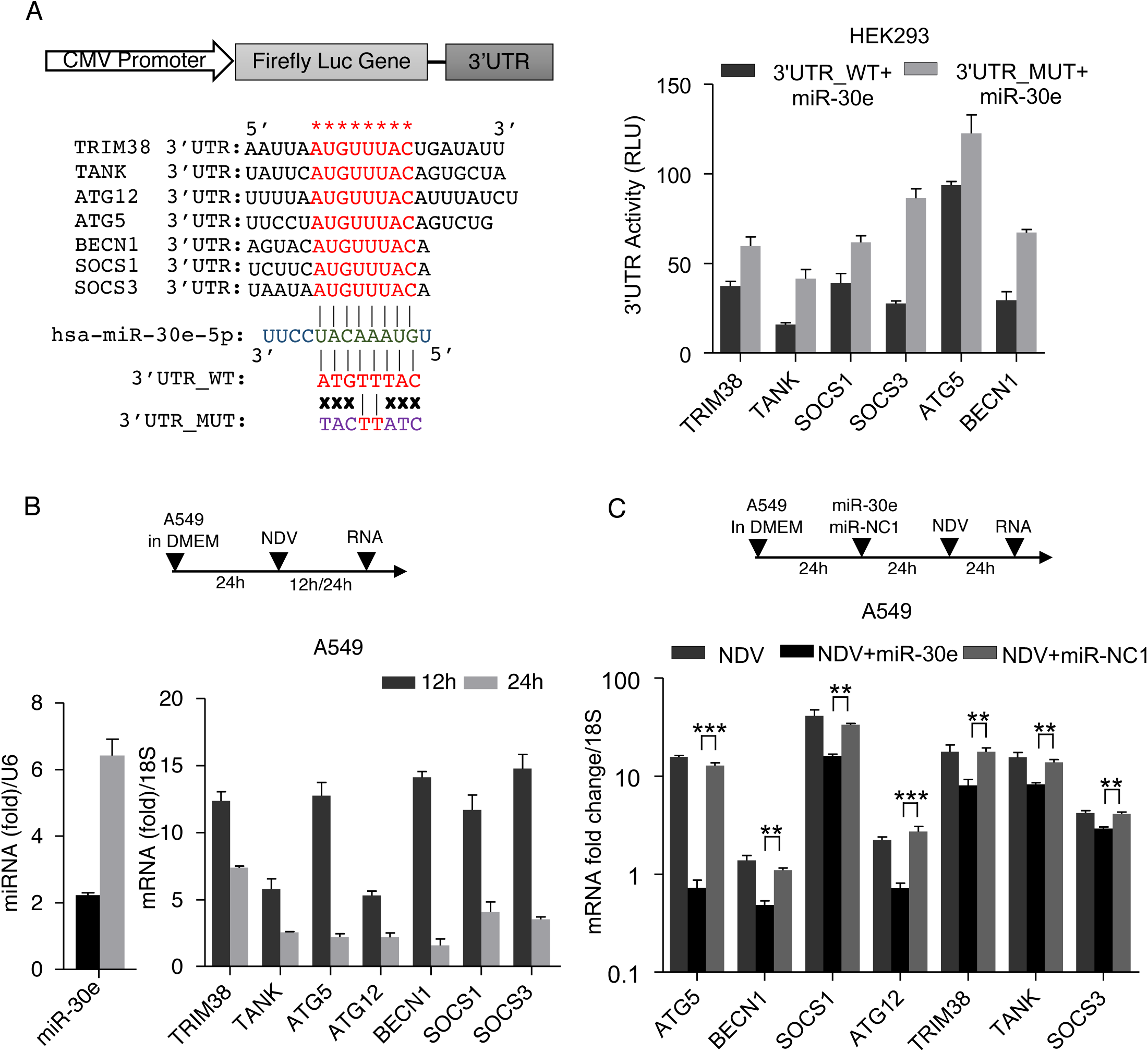
Quantification of innate immune negative regulators in presence of miR-30e: (A) HEK293 cells were transfected with miR-30e (25nM) mimic and 50 ng of pRL-TK along with 300ng of 3’UTR_WT or 300ng of 3’UTR_MUT for 24 hours, the cell was lysed and subjected to luciferase assay. 3’-UTRs of all selected target genes having binding sites for seed sequence in miR-30e were conserved throughout (shown in red). (B and C) Quantification of the fold changes by qRT-PCR analysis in the relative abundances of miR-30e (at the indicated times), and negative regulator transcripts (*TRIM38, TANK, ATG5, ATG12, BECN1, SOCS1* and *SOCS3*) after infection of (B) NDV (MOI = 5) in A549 cells and (C) NDV (MOI = 5) for 24 hours in A549 cells remain untransfected, transfected with miR-30e or miR-NC1 as indicated. Data are mean +/-SEM of triplicate samples from single experiment and are representative of two independent experiments. \*\*\**P<*0.001, \*\**P<*0.01 and \**P<*0.05 by one-way ANOVA Tukey test.

**Figure S10.**
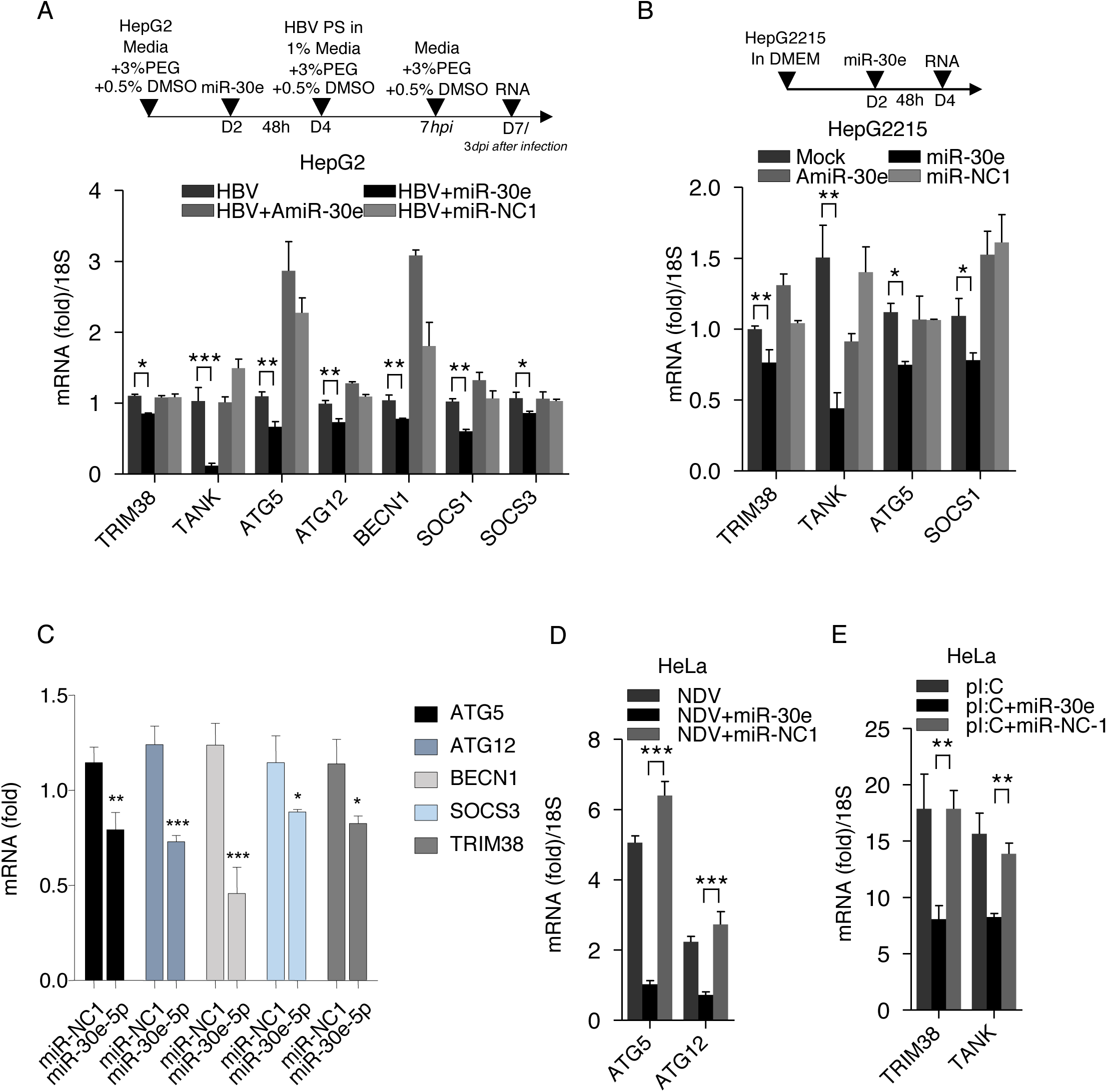
Quantification of innate immune negative regulators in presence of miR-30e: (A-D) Quantification of the fold changes by qRT-PCR analysis in the relative abundances of negative regulator transcripts (TRIM38, TANK, ATG5, ATG12, BECN1, SOCS1 and SOCS3) in (A) HepG2 cells remain untransfecd, transfected with miR-30e, AmiR-30e or miR-NC1 for 48 hours then treated with HBV-PS for HBV infection, (B) HepG2215 cells transfected as described previously, (C) HepG2-NTCP cells transfected with miR-30e and miR-NC1 for 48 hours then infected with HBV infection. (D-E) HeLa cells remain untransfecd, transfected with miR-30e, AmiR-30e or miR-NC1 for 24 hours then (D) infected with NDV (MOI = 5) or (E) treated with poly IC for 24 hours. D; represents number of days, dpi; days post infection and hpi; hours post infection. Data are mean +/-SEM of triplicate samples from single experiment and are representative of two independent experiments. ***P<0.001, **P<0.01 and *P<0.05 by one-way ANOVA Tukey test.

**Figure S11.**
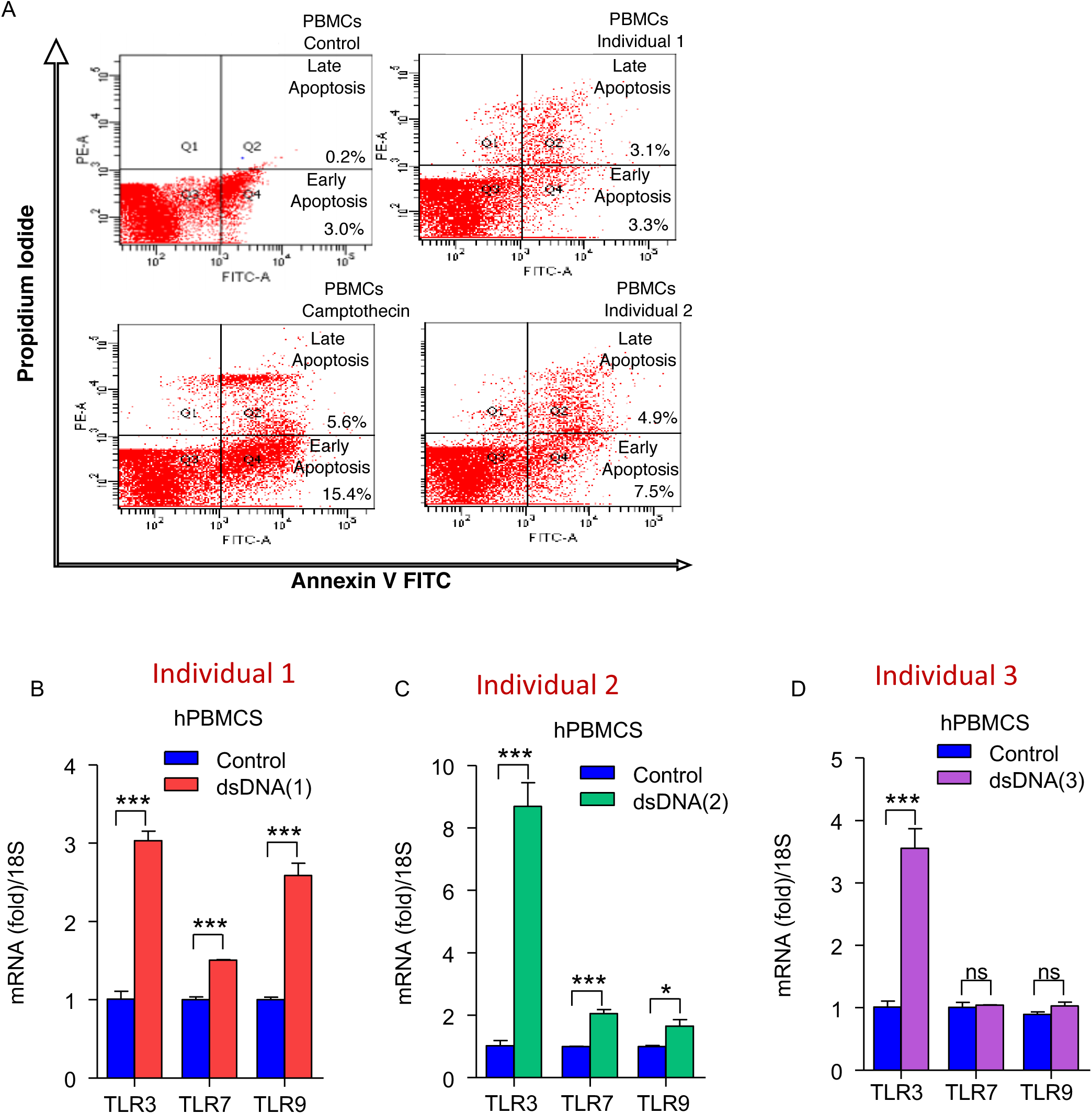
DAMPs induce apoptosis and TLR 3/7/9: (A) Human PBMCs remain untreated or treated with camptothecin, dsDNA separately from two different individuals (as indicated 1 and 2) and subjected to Annexin PI assay to detect the apoptosis level within the PBMCs using flow cytometery. (B-D) Quantification of the fold changes by qRT-PCR analysis in the relative abundances of respective transcripts (TLR3, TLR7 and TLR9) in all individuals. Data is the representative of three independent experiments. Data are mean +/-SEM of triplicate samples from single experiment and are representative of three independent experiments. ***P<0.001 and **P<0.01 by unpaired t-test.

**Figure S12.**
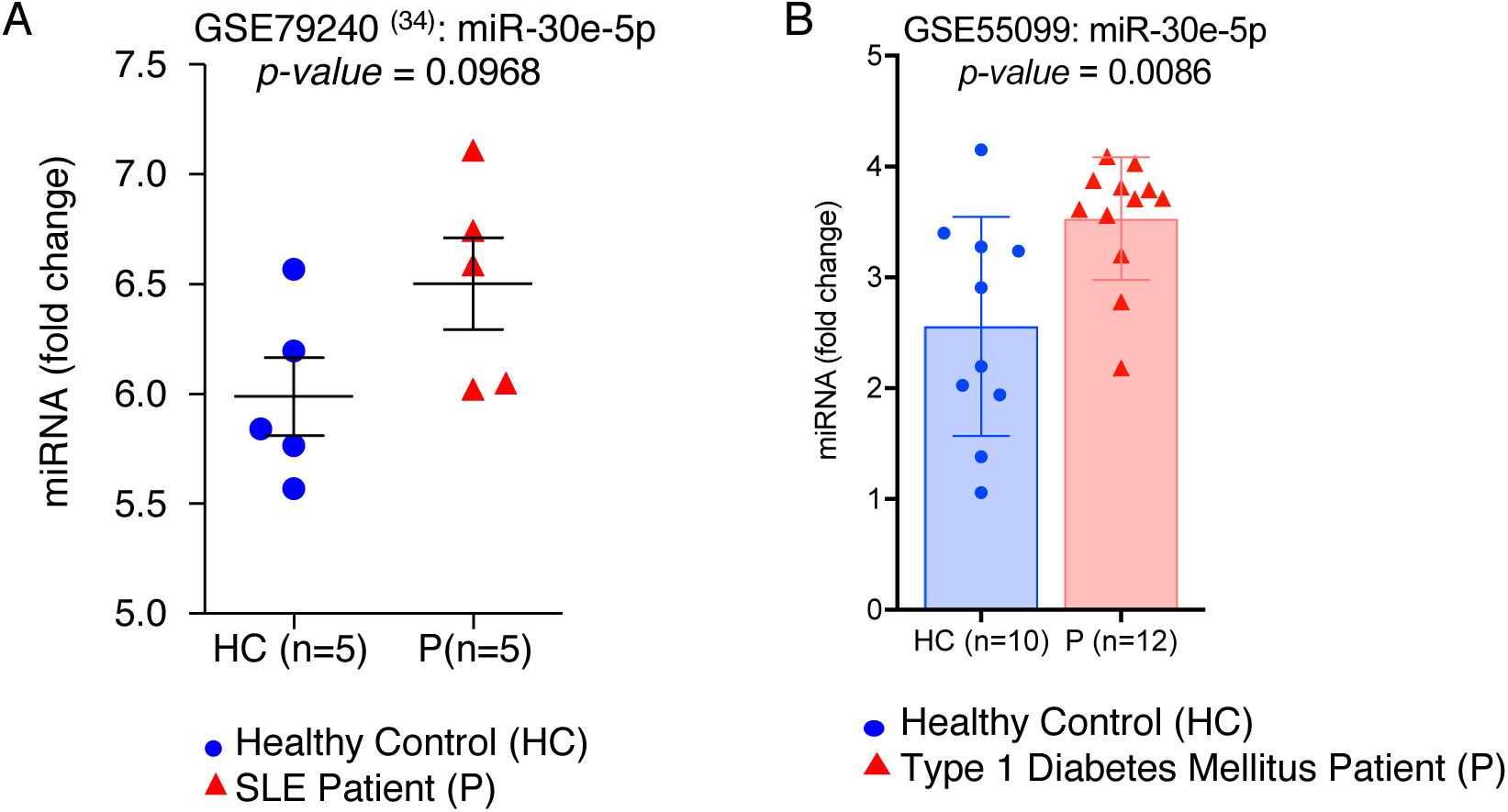
GEO Datasets re-analyzed to demonstrate the expression level of miR-30e in autoimmune disorder: (A) SLE (GSE79240) - Non-coding RNA profiling by microarray in dendritic cells of SLE patients (P) compared to healthy controls (HC). Data points include fold change of miR-30e among 5 patients. (B) Type 1 Diabetes Mellitus (GSE55099) - Non-coding RNA profiling by microarray in PBMCs of patients (P) compared to healthy controls (HC). Data points include fold change of miR-30e among 12 patients compared to 10 healthy controls (*P-value* = 0.0086).

**Figure S13.**
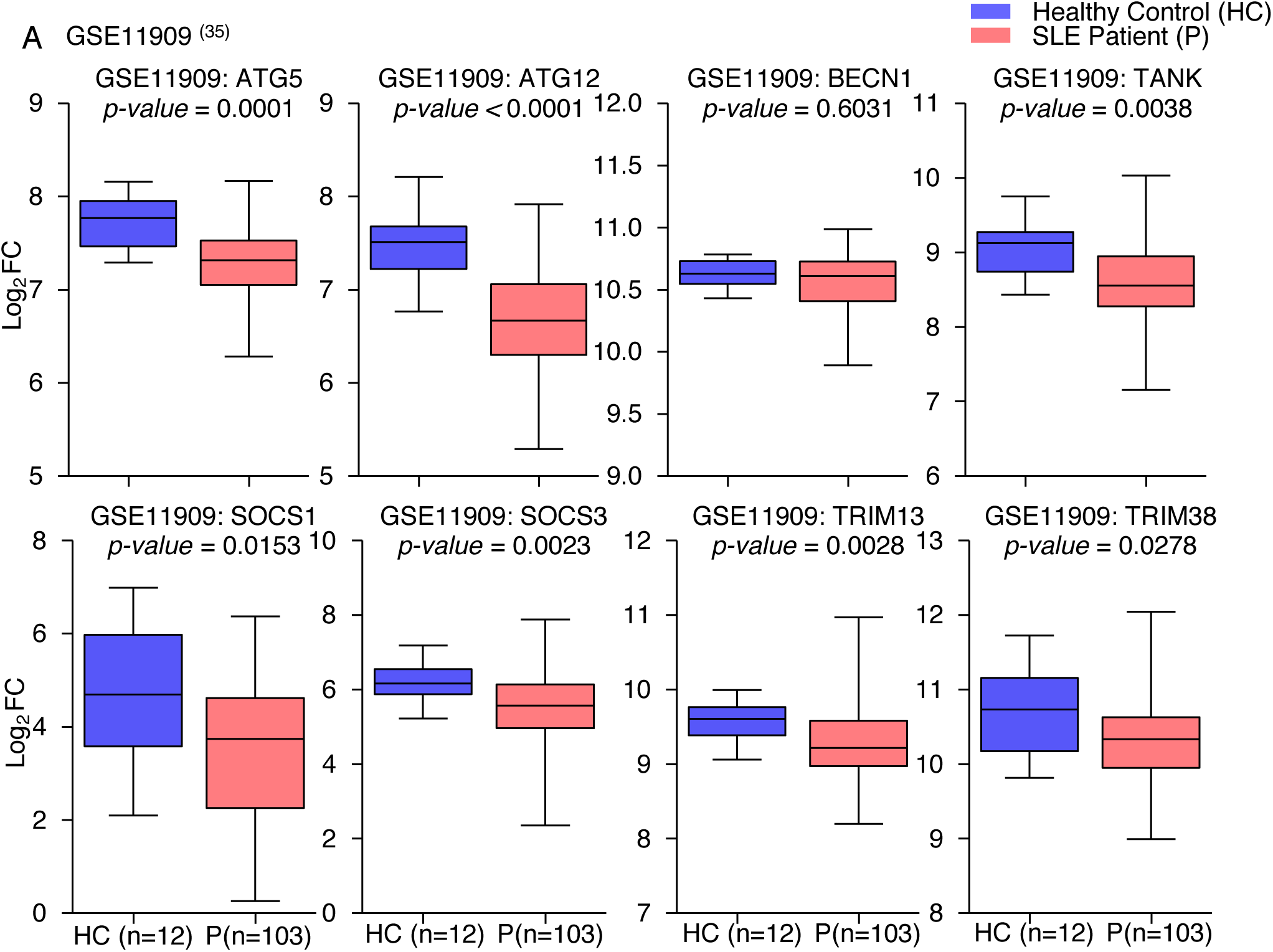
**– GEO Dataset-G**SE11909 **re-analyzed to demonstrate the transcript levels of innate negative regulators in SLE:** (A) Expression profiling by microarray to estimate the log fold change of following innate negative regulators during SLE pathogenesis in patients (P) compared to healthy controls (HC); ATG5, ATG12, BECN1, TANK, SOCS1, SOCS3, TRIM13 and TRIM38.

**Figure S14.**
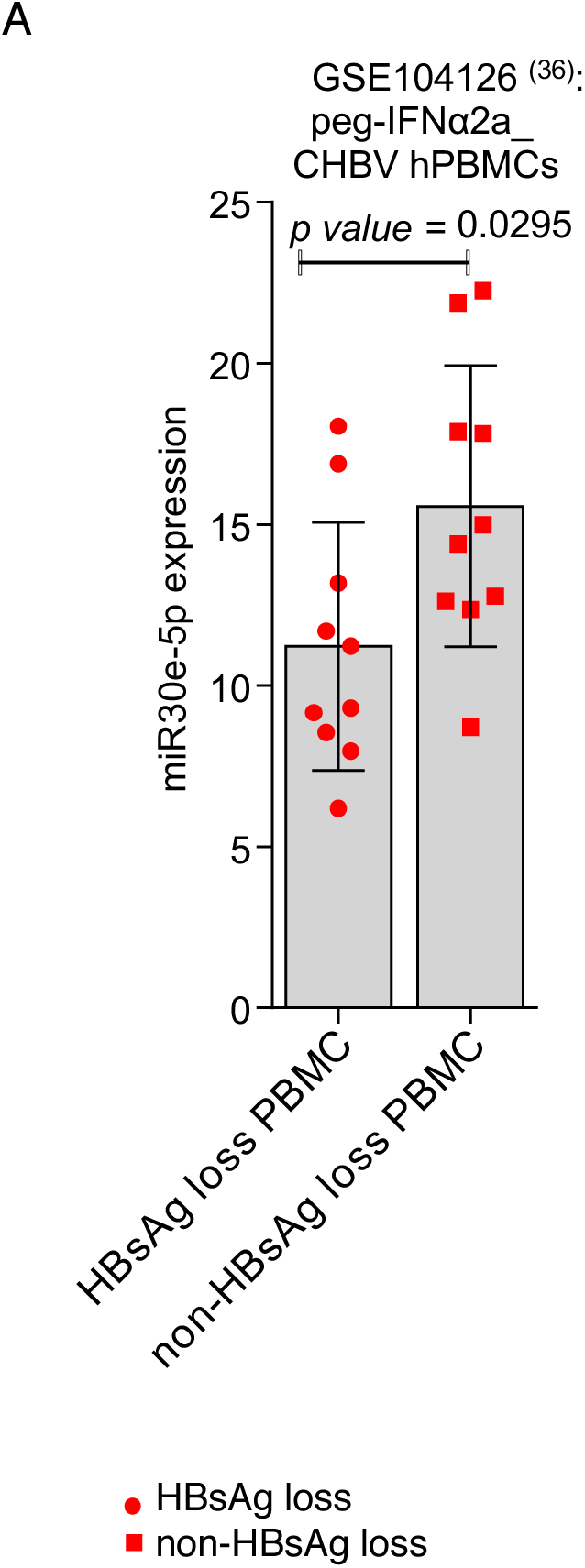
**– GEO Dataset-G**SE104126 **re-analyzed to demonstrate the expression level of miR-30e:** (A) Non-coding RNA profiling by microarray in PBMCs of two types of chronic hepatitis B (CHB) patients (HBsAg loss and non-HBsAg loss) after treatment with pegylated interferon (peg-IFN*α*2a). Data points include fold change of miR-30e among 10 patients.

## Supplementary Tables

**Table T1:** Demographic data of chronic hepatitis B (CHB) cohort and controls

|  | Therapy naive CHB<br>(n=51) | Control (n=24) | *p-value* |
| --- | --- | --- | --- |
| Median age (range) | 48(19-68) | 31(23-55) |  |
| Male/Female | 36/15 | 10/14 |  |
| AST<br>(reference range: 10 - 40 U/L) | 37(18-166) | 24(16-42) |  |
| ALT<br>(reference range:7-56U/L) | 36(15-130) | 23.5(12-48) |  |
| HBsAg | Positive | Negative |  |
| Anti-HBcIgM | Negative | ..... |  |
| HbeAg | 16 positive | ..... |  |
| HBV DNA Load log10<br>(IU/mL) | 5.03(1.36-9.2) | ..... |  |
| hsa-miR-30e-5p<br>(fold change/U6) | 18.2(0.63-269.12) | 1.01(0.712-1.92) | 0.0023*** |

Data expressed as median (range); CHB (Chronic Hepatitis B); AST (aspartate transaminase); ALT (alanine transaminase); HBsAg (Hepatitis B surface Antigen); Anti-HBcIgM (IgM antibody to hepatitis B core antigen); HBeAg (Hepatitis B e-Antigen); hsa-miR-30e-5p (Human Micro(mi)-RNA-30e).

**Table T2:**
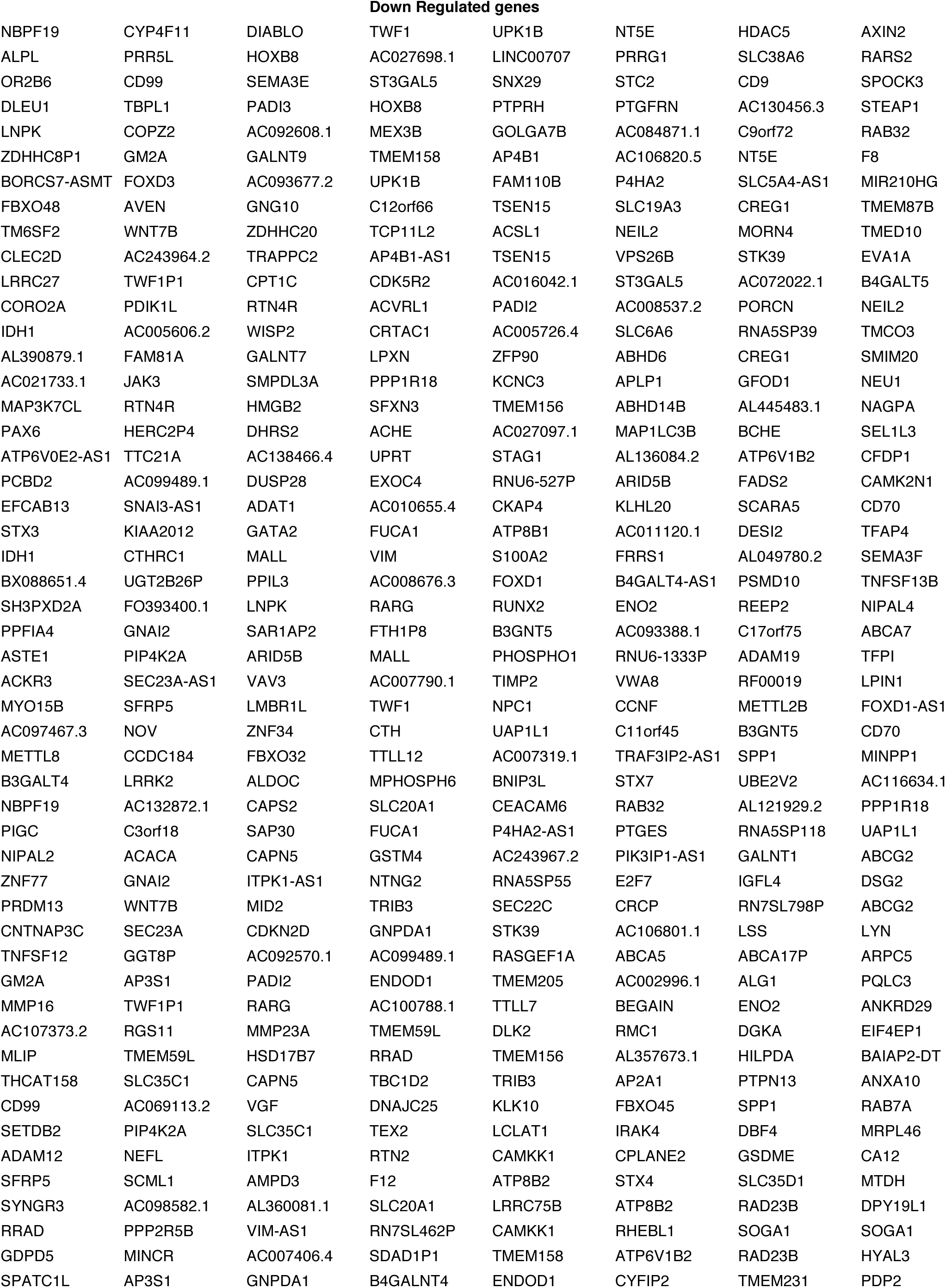
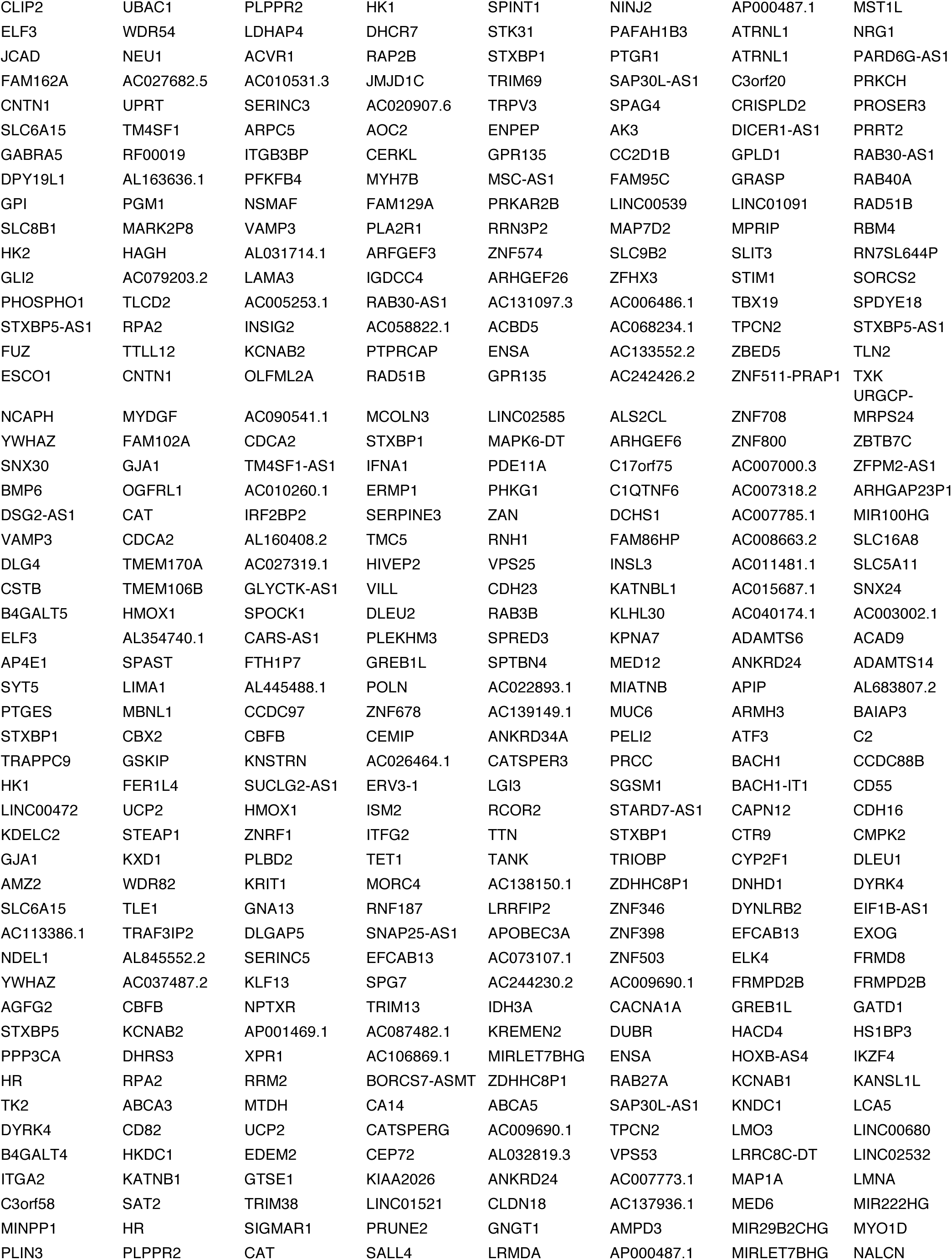
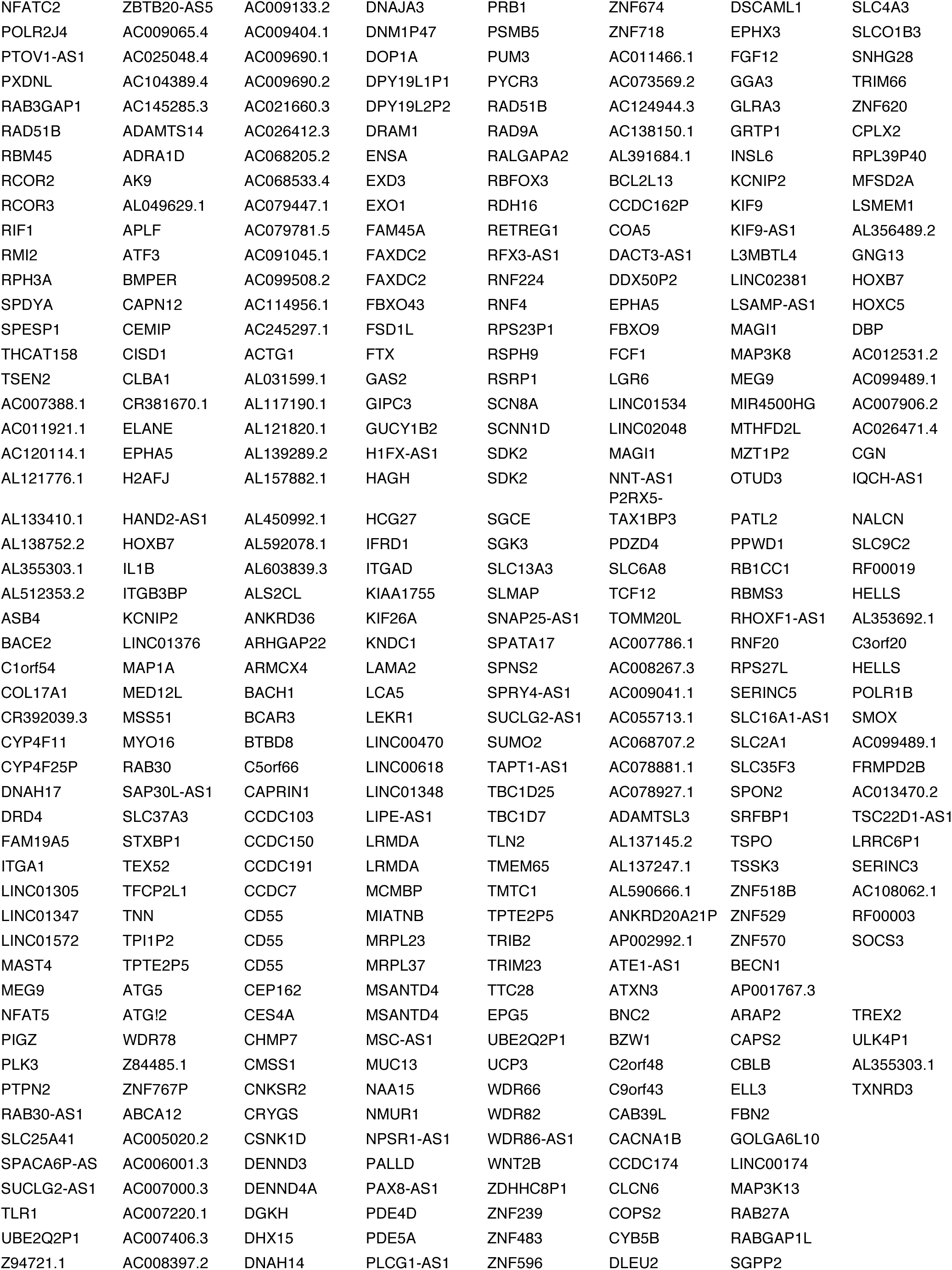
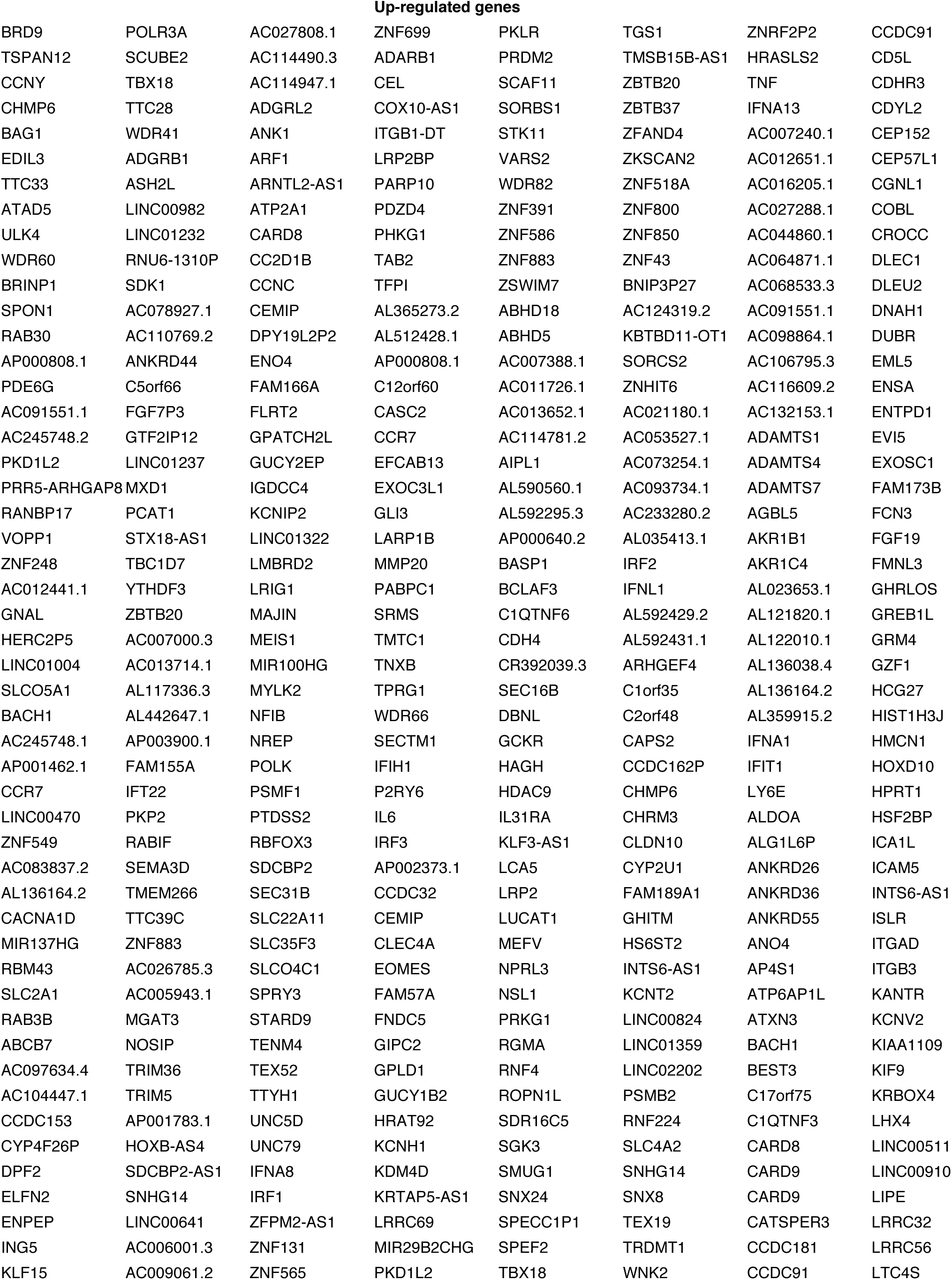
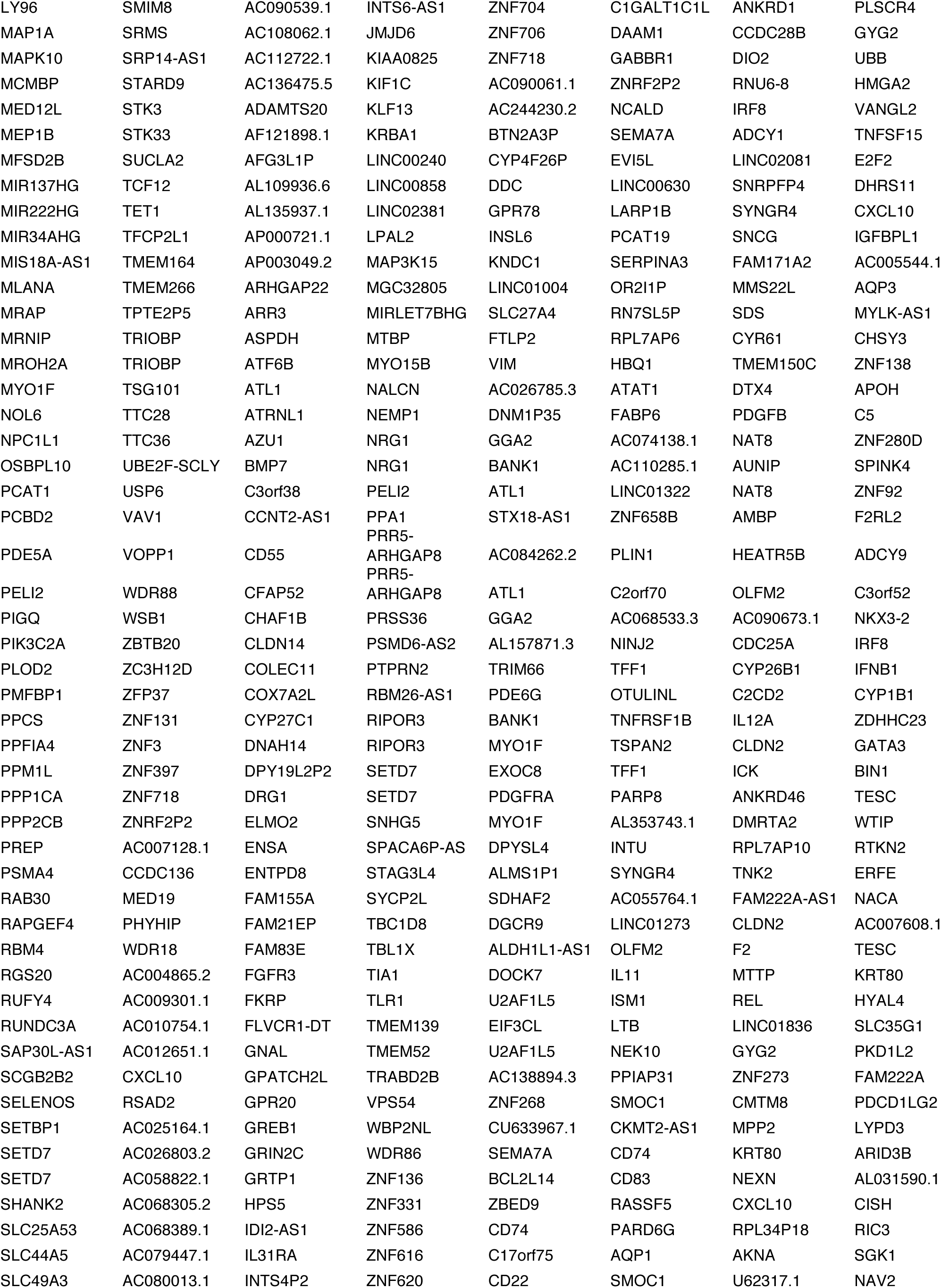
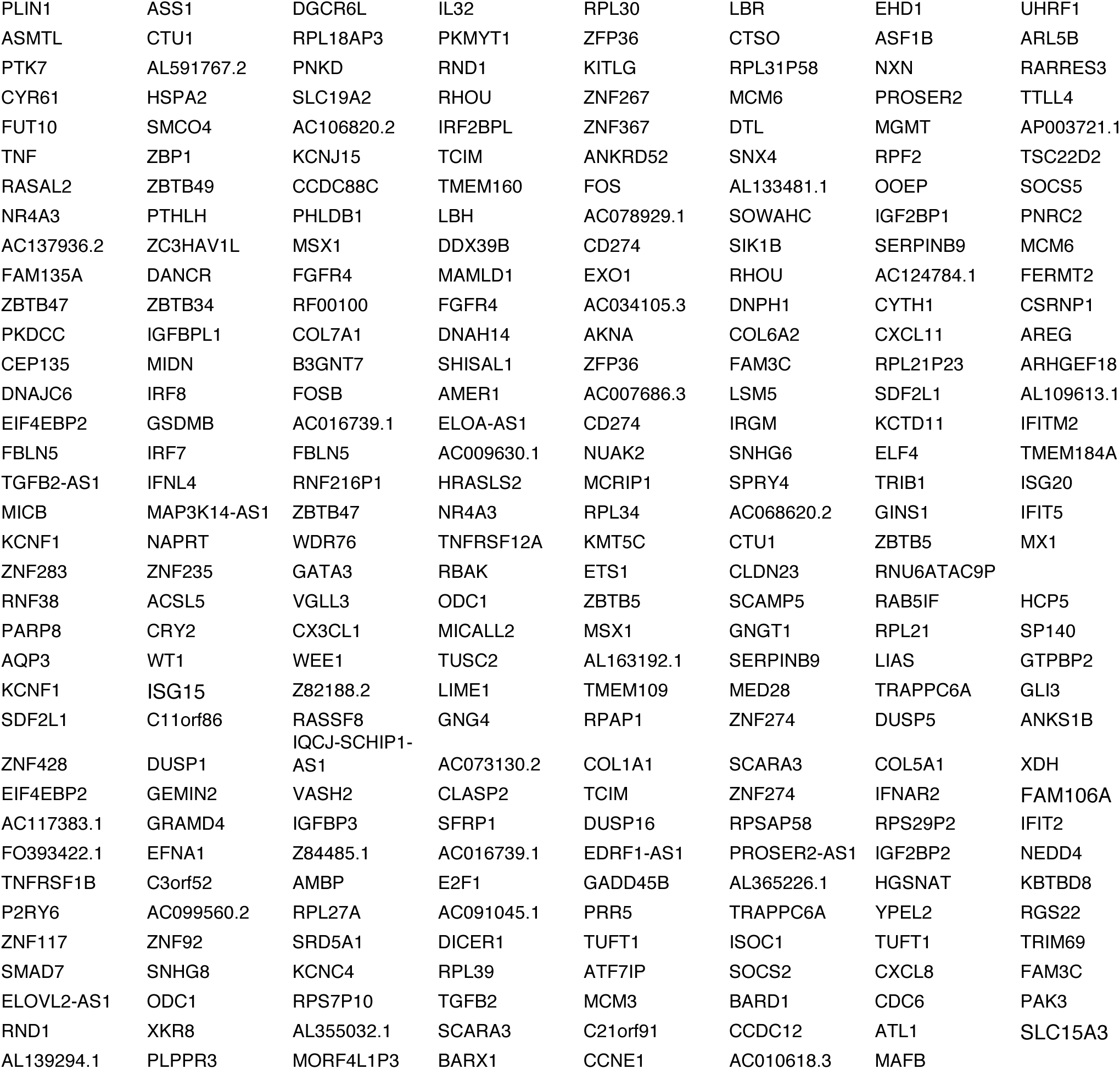
Differentially expressed genes by RNA sequencing analysis

**Table T3:**
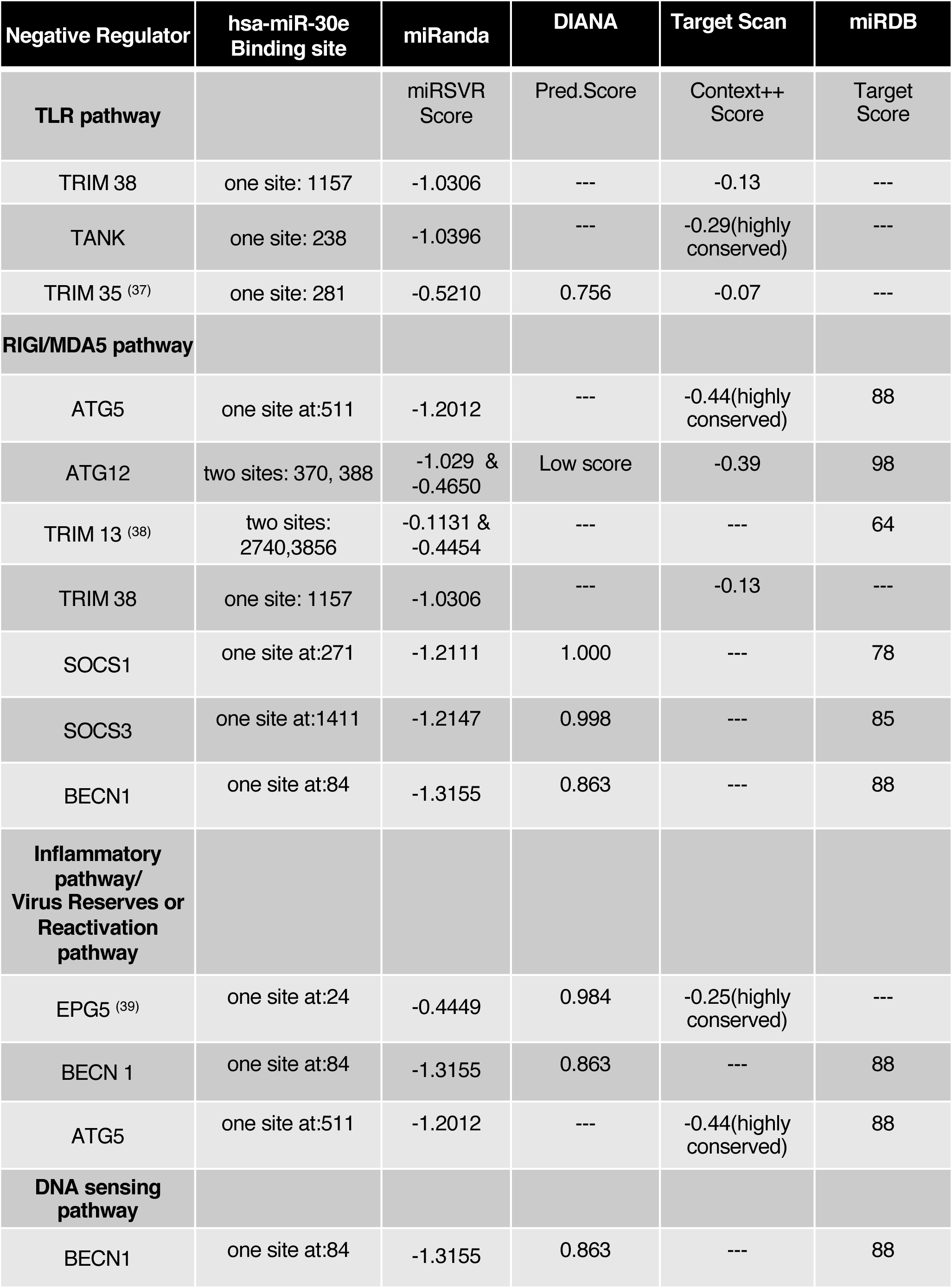
*In-silico* analysis of miR-30e targets using indicated algorithms

**Table T4:**
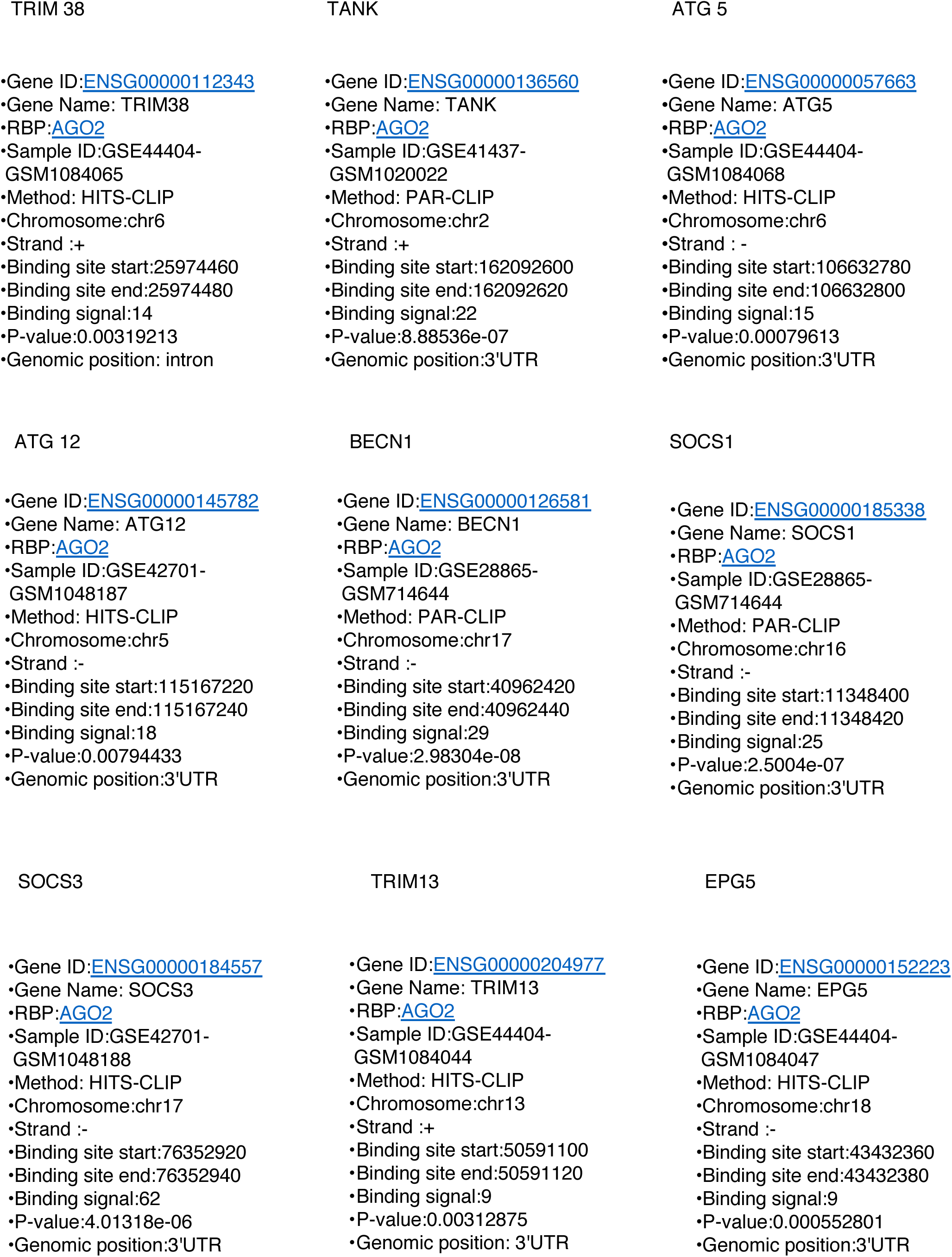
CLIP database analysis for miR-30e targets

**Table T5:**
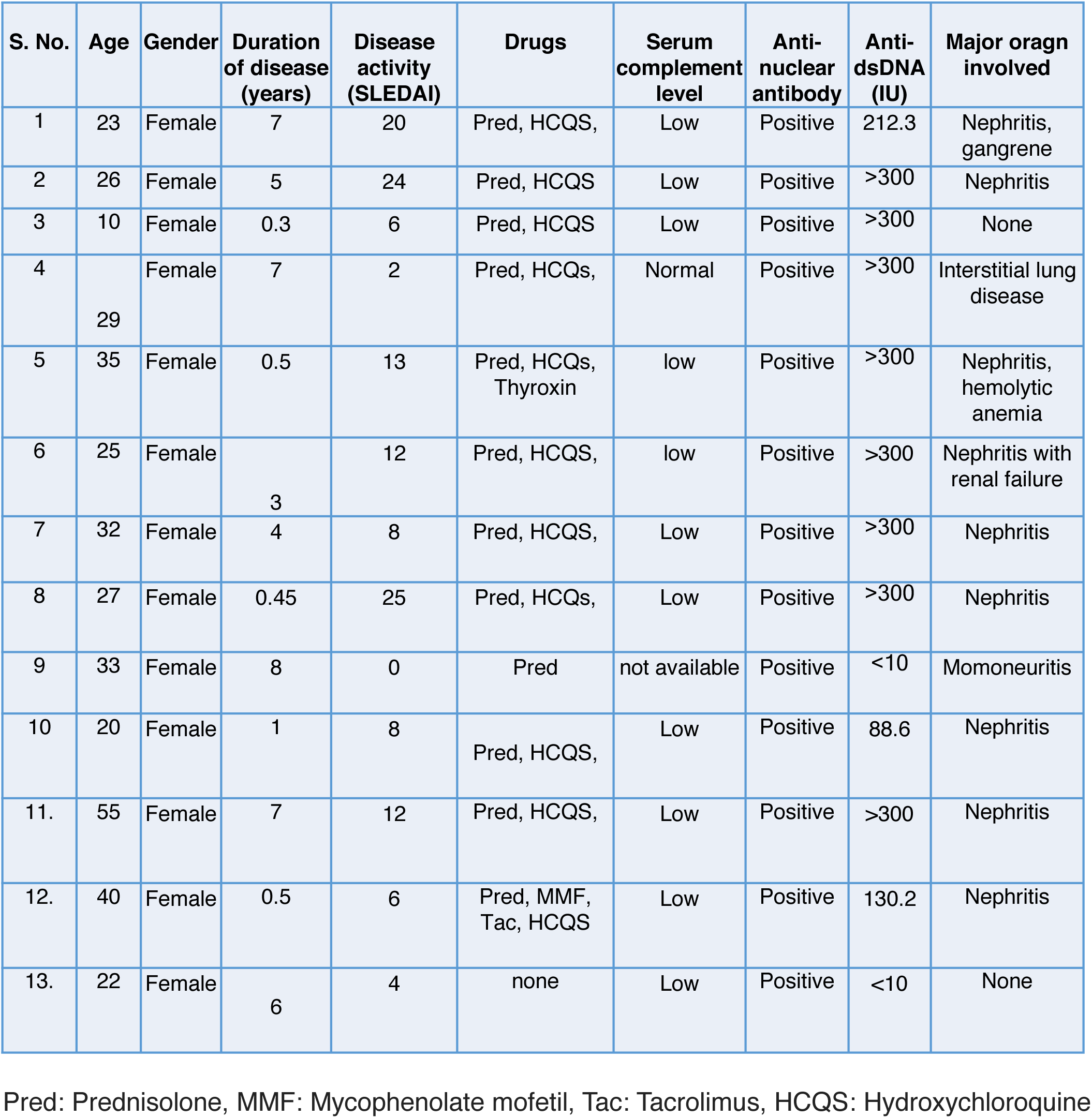
SLE patients details used in the study

**Table T6:**
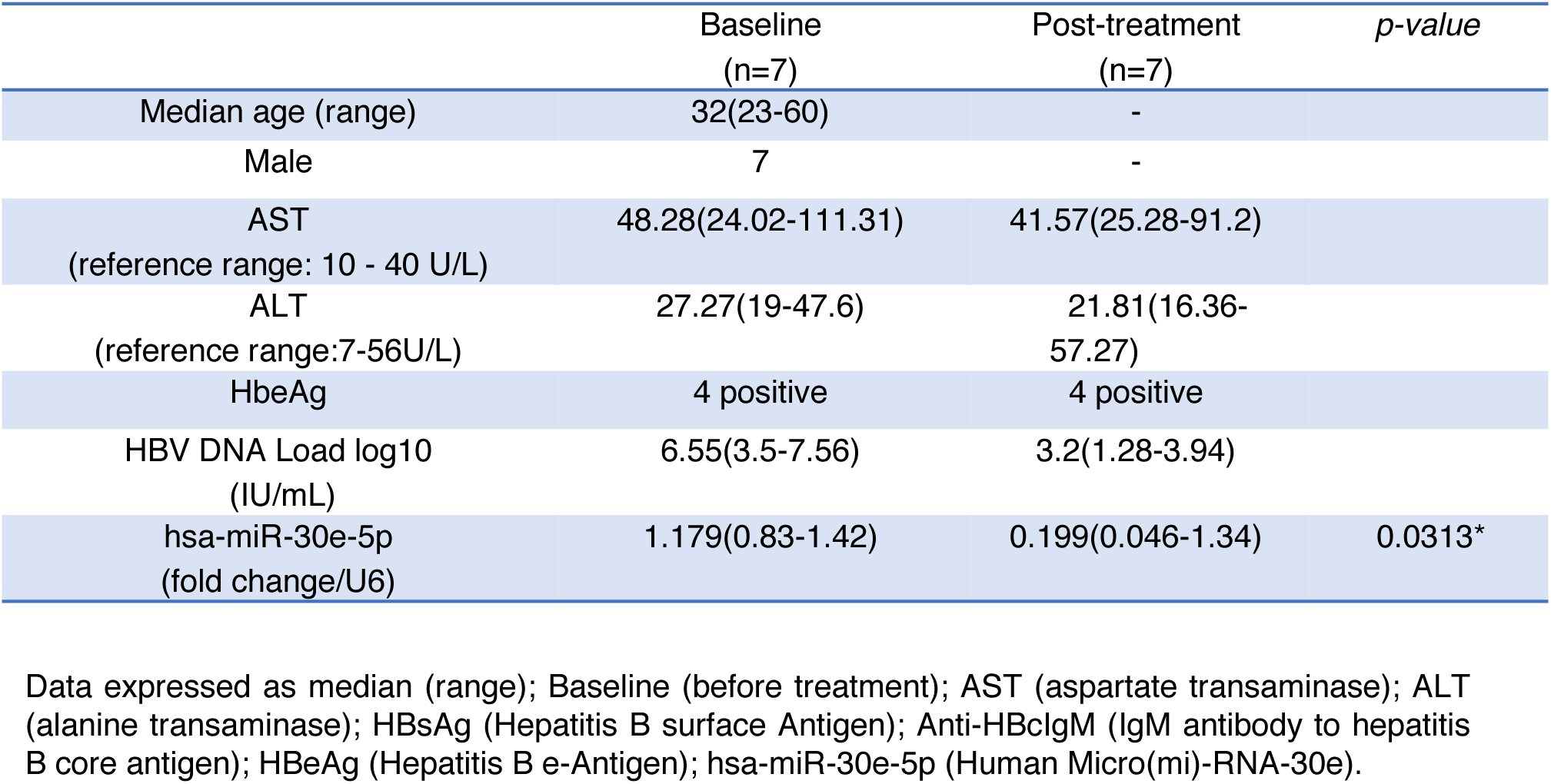
Demographic data of chronic hepatitis B (CHB) patients after Peg-interferon treatment

**Table T7:**
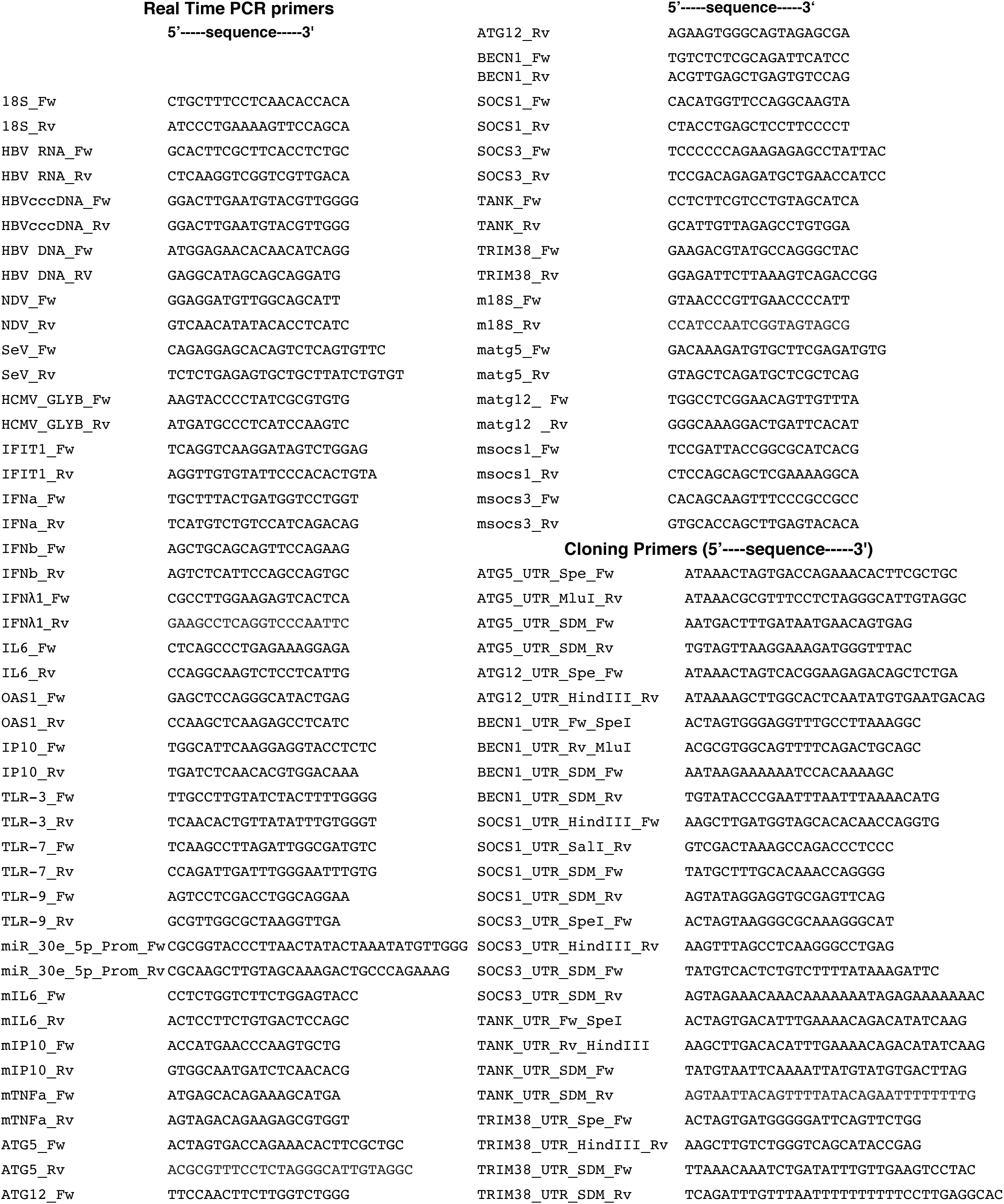
List of primers used in the study

